# Efficacy and breadth of adjuvanted SARS-CoV-2 receptor-binding domain nanoparticle vaccine in macaques

**DOI:** 10.1101/2021.04.09.439166

**Authors:** Hannah A. D. King, M. Gordon Joyce, Ines Elakhal Naouar, Aslaa Ahmed, Camila Macedo Cincotta, Caroline Subra, Kristina K. Peachman, Holly H. Hack, Rita E. Chen, Paul V. Thomas, Wei-Hung Chen, Rajeshwer S. Sankhala, Agnes Hajduczki, Elizabeth J. Martinez, Caroline E. Peterson, William C. Chang, Misook Choe, Clayton Smith, Jarrett A. Headley, Hanne A. Elyard, Anthony Cook, Alexander Anderson, Kathryn McGuckin Wuertz, Ming Dong, Isabella Swafford, James B. Case, Jeffrey R. Currier, Kerri G. Lal, Mihret F. Amare, Vincent Dussupt, Sebastian Molnar, Sharon P. Daye, Xiankun Zeng, Erica K. Barkei, Kendra Alfson, Hilary M. Staples, Ricardo Carrion, Shelly J. Krebs, Dominic Paquin-Proulx, Nicos Karasavvas, Victoria R. Polonis, Linda L. Jagodzinski, Sandhya Vasan, Paul T. Scott, Yaoxing Huang, Manoj S. Nair, David D. Ho, Natalia de Val, Michael S. Diamond, Mark G. Lewis, Mangala Rao, Gary R. Matyas, Gregory D. Gromowski, Sheila A. Peel, Nelson L. Michael, Kayvon Modjarrad, Diane L. Bolton

## Abstract

Emergence of novel variants of the severe acute respiratory syndrome coronavirus-2 (SARS-CoV-2) underscores the need for next-generation vaccines able to elicit broad and durable immunity. Here we report the evaluation of a ferritin nanoparticle vaccine displaying the receptor-binding domain of the SARS-CoV-2 spike protein (RFN) adjuvanted with Army Liposomal Formulation QS-21 (ALFQ). RFN vaccination of macaques using a two-dose regimen resulted in robust, predominantly Th1 CD4+ T cell responses and reciprocal peak mean neutralizing antibody titers of 14,000-21,000. Rapid control of viral replication was achieved in the upper and lower airways of animals after high-dose SARS-CoV-2 respiratory challenge, with undetectable replication within four days in 7 of 8 animals receiving 50 µg RFN. Cross-neutralization activity against SARS-CoV-2 variant B.1.351 decreased only ∼2-fold relative to USA-WA1. In addition, neutralizing, effector antibody and cellular responses targeted the heterotypic SARS-CoV-1, highlighting the broad immunogenicity of RFN-ALFQ for SARS-like betacoronavirus vaccine development.

**Significance Statement:** The emergence of SARS-CoV-2 variants of concern (VOC) that reduce the efficacy of current COVID-19 vaccines is a major threat to pandemic control. We evaluate a SARS-CoV-2 Spike receptor-binding domain ferritin nanoparticle protein vaccine (RFN) in a nonhuman primate challenge model that addresses the need for a next-generation, efficacious vaccine with increased pan-SARS breadth of coverage. RFN, adjuvanted with a liposomal-QS21 formulation (ALFQ), elicits humoral and cellular immune responses exceeding those of current vaccines in terms of breadth and potency and protects against high-dose respiratory tract challenge. Neutralization activity against the B.1.351 VOC within two-fold of wild-type virus and against SARS-CoV-1 indicate exceptional breadth. Our results support consideration of RFN for SARS-like betacoronavirus vaccine development.

## INTRODUCTION

The coronavirus infectious disease 2019 (COVID-19) pandemic, precipitated by severe acute respiratory syndrome coronavirus-2 (SARS-CoV-2), continues to threaten global public health and economies. Threats of future outbreaks also loom, as evidenced by three emergent SARS-like diseases caused by zoonotic betacoronaviruses in the last two decades. While several emergency use authorized (EUA) vaccines currently in use are expected to curb both disease and transmission of SARS-CoV-2 (1–6), the emergence of circulating variants of concern (VOC) less sensitive to vaccine-elicited immunity has raised concerns for sustained vaccine efficacy (7). Logistic challenges of vaccine production, distribution, storage and access for these vaccines will need to be resolved equitably to achieve resolution to the pandemic (8, 9). The rapid and unparalleled spread of SARS-CoV-2 has driven an urgent need to deploy scalable vaccine platforms to combat the ongoing pandemic and mitigate future outbreaks.

Current vaccines primarily focus the immune response to the spike glycoprotein (S) on the virion surface as it mediates host cell viral fusion and entry. The receptor-binding domain (RBD) of S engages the primary host cell receptor, Angiotensin-converting enzyme 2 (ACE2), for both SARS-CoV-2 and SARS-CoV-1, making RBD a promising domain for vaccine elicited immune focus (10–12). Moreover, many of the potently neutralizing monoclonal antibodies isolated against SARS-CoV-2 target the RBD (13, 14). Vaccination of nonhuman primates with RBD-encoding RNA or DNA protects against respiratory tract challenge, indicating that immune responses to the RBD can prevent viral replication (15, 16). RBD vaccination also elicits cross-reactive responses to circulating SARS-CoV-2 VOC in both animals and humans (17, 18), with decrements against the more difficult to neutralize B.1.351 variant similar to that seen with S immunogens (19). The breadth of RBD immunogenicity is further supported by the ability of RBD-specific monoclonal antibodies isolated from SARS-CoV-1 convalescent individuals to cross-neutralize SARS-CoV-2 (20, 21). These findings suggest potential for RBD-based vaccines being efficacious against SARS-CoV-2 variants and other coronavirus species.

Approaches to improve immunogenicity of S or RBD protein vaccines include optimizing antigen presentation and co-formulating with adjuvants to enhance the protective immunity. One common approach to enhance the elicitation of adaptive immune responses is the multimeric presentation of antigen, for example, on the surface of nanoparticles or virus-like particles (22). Presenting RBD in ordered, multivalent arrays on the surface of self-assembling protein nanoparticles is immunogenic and efficacious in animals (23–28), with improved immunogenicity relative to monomeric soluble RBD and cross-reactive responses to variants (17, 24, 26). However, it is unknown whether RBD nanoparticle vaccines are able to protect against infection in primates, which have become a standard model for benchmarking performance of vaccine candidates. Liposomal adjuvants incorporating QS-21, such as that used in the efficacious varicella zoster vaccine, SHINGRIX^®^, may augment protective immunity to SARS-CoV-2 vaccines. Such adjuvants have previously demonstrated superior humoral and cellular immunogenicity relative to conventional adjuvants (29, 30).

Here, we evaluate the use of a ferritin nanoparticle vaccine presenting the SARS-CoV-2 RBD (RFN) adjuvanted with the Army Liposomal Formulation QS-21 (ALFQ) (31). Both ferritin nanoparticles and ALFQ have been evaluated for vaccination against multiple pathogens in humans in phase 1 clinical trials (32–34). We demonstrate in a nonhuman primate model that immunization with RFN induces robust and broad antibody and T cell responses, as well as protection against viral replication and lung pathology following high-dose respiratory tract challenge.

## RESULTS

### Vaccine and animal study design

A SARS-CoV-2 RBD ferritin nanoparticle vaccine (RFN) was designed as a ferritin-fusion recombinant protein that self-assembles into a 24-mer nanoparticle displaying a multivalent, ordered array of RBD on its surface. Briefly, the RBD protein sequence (residues 331-527) derived from the Wuhan-Hu-1 genome sequence (GenBank accession number: MN908947.3) was covalently linked to the C-terminal region of the *Helicobacter pylori* ferritin molecule. Twenty-three rhesus macaques were immunized with either 50 or 5 μg of RFN, or sham-immunized with PBS (n=7/8 per group), at study weeks 0 and 4 (Fig. 1*A*). RFN was adjuvanted with ALFQ, which contains synthetic monophosphoryl 3-deacyl lipid A and QS-21. Animals were challenged 4 weeks after the last immunization via combined intratracheal (IT, 1.0 mL) and intranasal (IN, 0.5 mL per nostril) inoculation of a 10^6^ TCID50 dose of SARS-CoV-2 virus (USA-WA1/2020). Animals were followed for 7 (n=12) or 14 days (n=11) following challenge for immunologic, virologic, and pathologic assessments.

**Figure 1.**
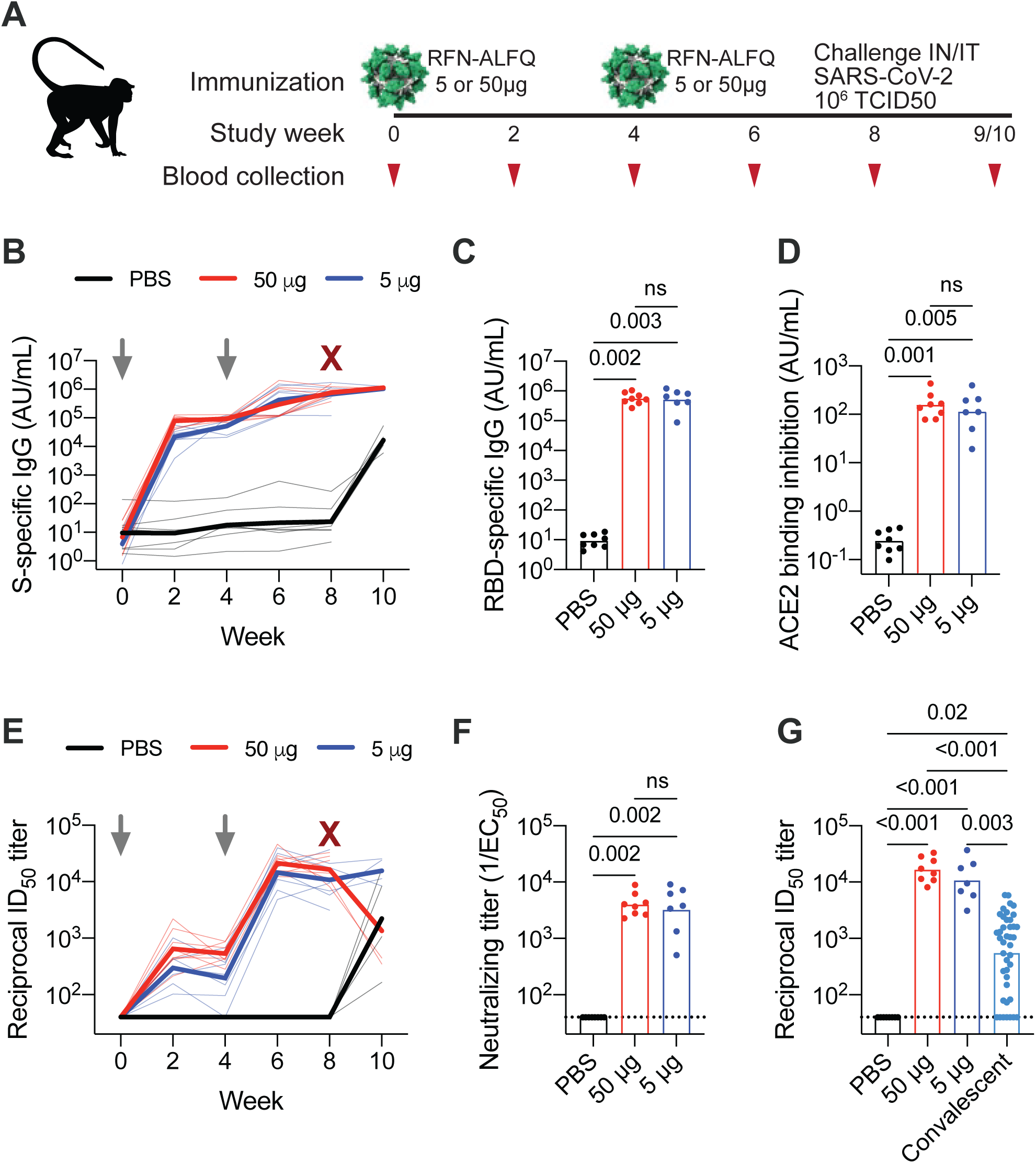
RFN vaccine-elicited binding and neutralizing antibody responses to SARS-CoV-2. Humoral immune responses were measured in vaccinated macaques. (*A*) Rhesus macaque vaccination, challenge, and sampling schedule. Animals were immunized with either 50 or 5 µg RFN-ALFQ at weeks 0 and 4; control animals received PBS (N= 7-8 per group). 1×10^6^ TCID50 of SARS-CoV-2 was administered four weeks after the last vaccination. *(B)* Serum SARS-CoV-2 S-specific IgG responses assessed by MSD every two weeks following vaccination. Data are depicted as the arbitrary units (AU)/ml of IgG binding. Thick lines indicate geometric means within each group and thin lines represent individual animals. Serum SARS-CoV-2 RBD-specific IgG *(C)* and inhibition of angiotensin-converting enzyme 2 (ACE2) binding to the RBD *(D)* four weeks after last vaccination were measured by MSD. *(E)* Serum SARS-CoV-2 S-specific pseudovirus neutralization every two weeks following vaccination. Virus neutralization reciprocal 50% inhibitory dilution (ID50) is shown. Thick lines indicate geometric means within each group and thin lines represent individual animals. *(F)* Authentic SARS-CoV-2 WA1 virus neutralization at four weeks after last vaccination. *(G)* Pseudovirus neutralization activity four weeks post-boost was compared to a panel of human convalescent sera (N=41 samples). Bars indicate the geometric mean titer. Symbols represent individual animals and overlap with one another for equal values where constrained. Gray arrows indicate the time of immunization, maroon arrow indicates time of challenge. Significance was assessed using a Kruskal-Wallis test followed by a Dunn’s post-test.

### Humoral responses to vaccination

Multiple vaccine-matched humoral immune responses were measured longitudinally in serum following vaccination. First, binding antibody responses to the SARS-CoV-2 prefusion stabilized S protein (S-2P) (35) were assessed by MSD. Immunization with either 5 or 50 μg of RFN elicited S-specific IgG two weeks following the prime (21,896 and 79,109 AU/mL, respectively) (Fig. 1*B*). These responses increased two weeks following the second immunization (420,249 and 293,509 AU/mL). Boosting was greater with the 5 μg dose, achieving a 19-fold increase relative to post-prime versus ∼3.7-fold with 50 μg. Responses continued to marginally increase four weeks following the second immunization as well as two weeks post-challenge. Unvaccinated control animals mounted responses ∼1,000-fold over baseline within two weeks post challenge and these responses were ∼65-fold lower than those in vaccinated animals after challenge.

Given the importance of the RBD in mediating viral entry and the majority of neutralizing antibody responses targeting this domain, RBD-specific humoral responses were also measured. RFN induced binding antibodies four weeks following the second immunization, with no significant difference between vaccine dose groups (Fig. 1*C*). Responses in vaccinated animals were ∼60,000-fold over the background in unvaccinated controls and comparable in magnitude to those against the S protein, consistent with an RBD-focused response. To confirm these findings, the on-rate association between serum antibodies and RBD antigen was measured by biolayer interferometry and longitudinal responses tracked with S-binding and pseudovirus neutralizing responses (Fig. S1). Again, vaccine dose groups did not differ. Functional activity of serum antibodies to inhibit ACE2 binding to the RBD antigen was also measured and present, with high magnitude responses elicited by RFN at both the 5 and 50 μg doses (Fig. 1*D*).

Neutralizing antibody responses against SARS-CoV-2 using a pseudovirus neutralization assay followed a similar pattern as the S-specific binding responses (Fig. 1*E*). Peak ID50 titers of 14,540 and 21,298 were observed two weeks following the boost for the 5 and 50 μg RFN doses, respectively. Neutralizing responses increased markedly between the prime and boost, rising 48- and 32-fold between study weeks two and six. Among the 50 μg RFN vaccinated animals followed two weeks post-challenge, neutralizing responses declined six weeks post-boost by approximately one log relative to peak values, indicating neutralizing responses may decay more quickly than binding antibodies.

Neutralizing responses were also evaluated using an authentic SARS-CoV-2 virus neutralization assay (USA-WA1 isolate). Robust neutralizing titers were detected in all RFN vaccinated animals (Fig. 1*F*). Median EC50 titers were ∼3,800 for both dose groups, though slightly more variable with 5 μg dosing. This result paralleled responses assessed by a pseudovirus assay (Fig. 1*E,G*). Since serum from convalescent COVID-19 human cases is frequently used as a benchmarking reference for vaccine immunogenicity in clinical and pre-clinical studies, we compared RFN-vaccinated macaque pseudovirus neutralizing titers to those of 41 convalescent individuals 4-8 weeks post-COVID infection. Responses in the 50 μg group were on average 13-fold higher than the convalescent individuals, indicating that RFN elicited higher antibody titers than observed in the first months following human infection. Summarizing, RFN vaccination generated strong RBD-specific binding antibodies with potent neutralization activity which block the interaction between the RBD region of SARS-CoV-2 S and the host ACE2 receptor.

Non-neutralizing antibody effector functions associated with vaccine-mediated protection against other viruses may also be important for SARS-CoV-2 (36, 37). Strong IgG-mediated cellular opsonization responses were observed following the second immunization, while IgM and IgA were more modest (Fig. S2*A-C*). Serum antibody-dependent phagocytosis mediated by either monocytes (ADCP) or neutrophils (ADNP) as well as complement deposition (ADCD) responses were also robust in both vaccinated groups and consistently peaked at week six (Fig. S2*D-F*). A similar pattern was seen for antibody-dependent trogocytosis (38) (Fig. S2*G*). Overall, 5 µg RFN achieved equal Fc-mediated effector functions compared to 50 µg, though ADCD responses trended ∼1.25-fold greater with the higher dose.

### Virus-specific T cell responses

SARS-CoV-2-specific T cell immunity is associated with reduced disease severity and can influence antibody responses (39, 40). We assessed S-specific T cells in PBMCs by *in vitro* peptide stimulation and intracellular cytokine staining using a 19-color multiparameter flow cytometry panel for detailed functional characterization of T cell responses from RFN vaccination. A vigorous, dose-dependent Th1 (TNF, IL-2, IFN-*γ*) CD4+ T cell response was observed in all RFN vaccinated animals 4 weeks after the second vaccination, ranging from 0.4-5.2% of memory cells (Fig. 2*A*). These S-specific Th1 cells were polyfunctional in quality, a property associated with control of other pathogens (41), as a large proportion simultaneously expressed multiple Th1 cytokines (Fig. S3*A*). Th2 responses were low or undetectable (Fig. 2*B*), with median Th1/Th2 ratios of ∼20 among 50 µg-vaccinated animals with evidence of a Th2 response (Fig. S3*B*). CD8+ T cell responses were observed in about half of the animals and were more prominent in recipients of 50 μg than 5 μg RFN (Fig. S3*C*). Response magnitude was ∼0.1-0.4% of memory CD8+ T cells.

**Figure 2.**
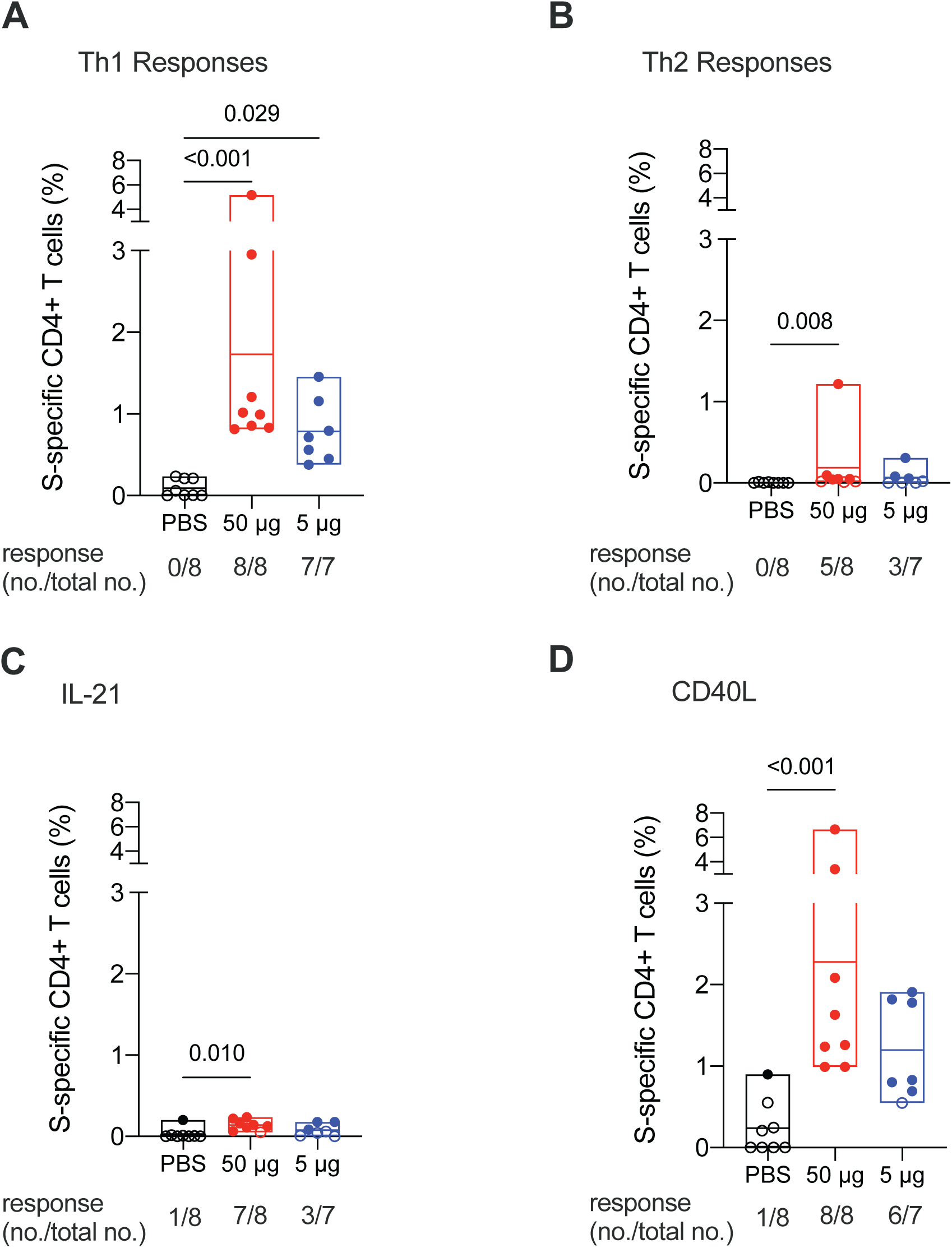
RFN vaccine elicited SARS-CoV-2 S-specific CD4+ T cell responses. T cell responses were assessed by SARS-CoV-2 S peptide pool stimulation and intracellular cytokine staining of peripheral blood mononuclear cells collected four weeks after last vaccination. S-specific memory CD4+ T cells expressing the indicated markers are shown as follows: (*A*) Th1 cytokines (IFN*γ*, TNF and IL-2); *(B)* Th2 cytokines (IL-4 and IL-13); *(C)* IL-21; *(D)* and CD40L. Boolean combinations of cytokine-positive memory CD4+ T cells were summed. Probable positive responses, defined as >3 times the group background at baseline, are depicted as closed symbols. Positivity rates within each group are shown below each graph as a fraction. Box plot horizontal lines indicate the mean; top and bottom reflect the minimum and maximum. Significance was assessed using a Kruskal-Wallis test followed by a Dunn’s post-test.

Additional CD4+ T cell functions important for the development of antibody responses were evaluated. S-specific CD4+ T cell IL-21 responses, a surrogate marker of peripheral T follicular helper cell activity, were observed in the majority of animals vaccinated with 50 µg and in half of the animals vaccinated with 5 µg RFN (Fig. 2*C*). The average frequency in responders was 0.15%. The CD4+ T cell activation marker, CD40L, which promotes B cell antibody isotype switching, was highly expressed by S-specific cells (Fig. 2*D*). Responses ranged from ∼1-7% after 50 μg RFN and were observed in all eight animals, while response rates and magnitude were slightly reduced with the 5 µg dose (∼0.7-2% in six of seven animals). Overall, these data show that adjuvanted RFN induced robust Th1-polarized polyfunctional CD4+ T cells favorable for viral clearance and with critical B-cell help capability.

### SARS-CoV-2 challenge efficacy

To assess the protective efficacy of RFN vaccination, animals were challenged with high-dose (10^6^ TCID50) SARS-CoV-2 USA-WA1 administered via the simultaneous IN/IT routes four weeks following the second immunization. The presence of viral RNA was assessed in both the upper (NP swabs and saliva) and lower (BAL) respiratory tract. Measurements were made of both total RNA and subgenomic E mRNA (sgmRNA), considered a more specific indicator of viral replication (42, 43). Unvaccinated control animals all showed evidence of a robust infection, with mean levels of sgmRNA in the BAL of ∼10^6^ copies/mL, and in the NP swabs of ∼10^7^ copies/mL at day 2 post-challenge (Fig. 3). Moreover, viral replication was sustained at >10^4^ copies/mL sgmRNA for 7 days in the upper airways. In RFN vaccinated animals, the magnitude and duration of viral replication was markedly reduced. In the 50 μg group, day 1 sgmRNA was reduced by 1 and 2 logs in the BAL and NP swabs, respectively. Rapid clearance was observed by day 2 in five of eight animals in the upper airways and four of eight in the lower airways. Both airways were void of replicating virus in all but one animal by day 4. Viral control was also apparent after 5 μg RFN vaccination, though with slightly more breakthrough replication early after challenge. The majority of animals had no detectable sgmRNA by day 4 in both the upper and lower airways.

**Figure 3.**
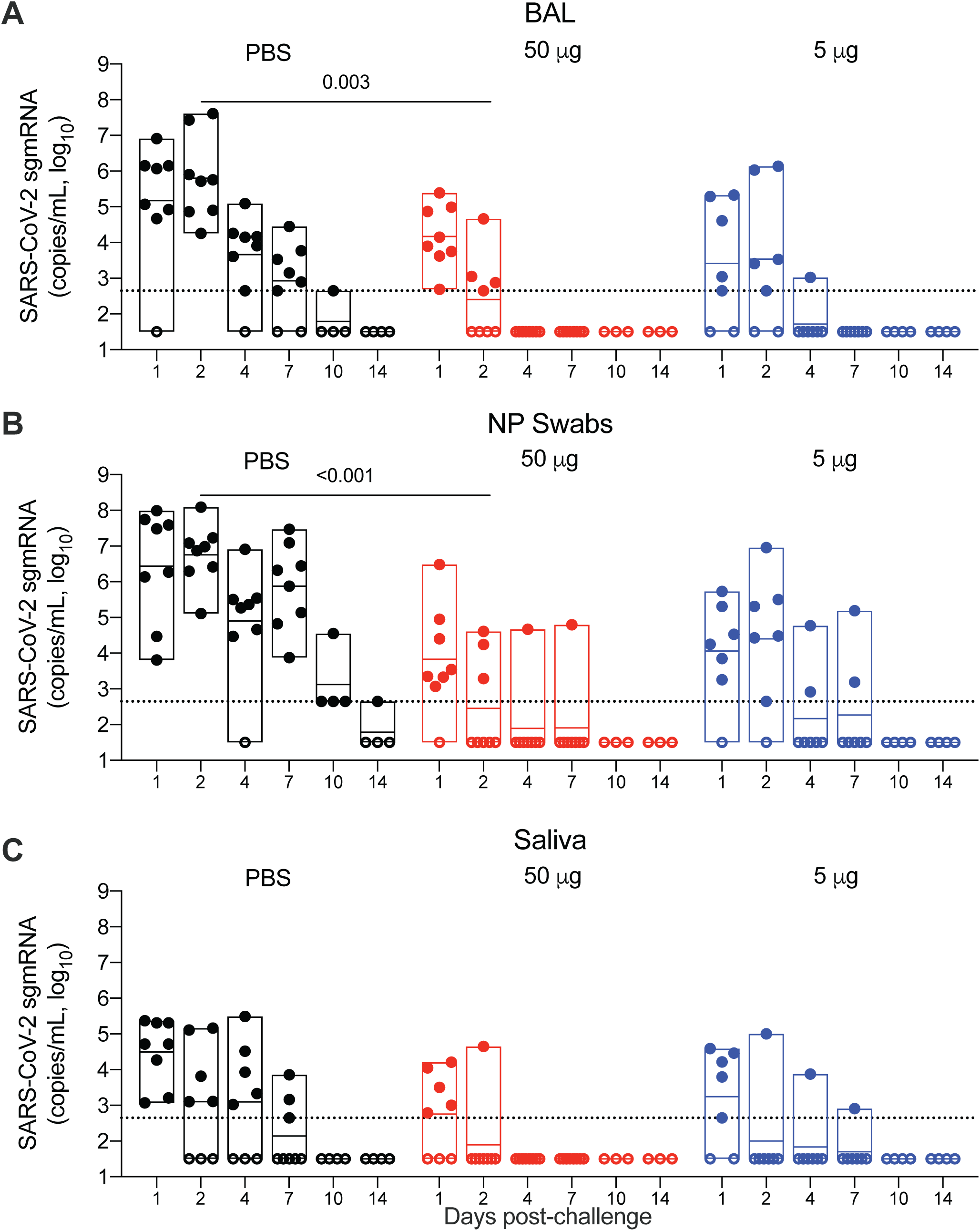
Viral replication in the lower and upper airways after RFN vaccination and subsequent SARS-CoV-2 respiratory challenge. Subgenomic E messenger RNA (sgmRNA) copies per milliliter were measured following challenge. *(A)* Bronchoalveolar lavage fluid (BAL), *(B)* nasopharyngeal (NP) swabs and *(C)* saliva of vaccinated and control animals for one (N=7-8 per group) or two weeks (N=3-4 per group) following intranasal and intratracheal SARS-CoV-2 (USA-WA1/2020) challenge of vaccinated and control animals. Specimens were collected 1, 2, 4, 7, 10 and 14 days post-challenge. Dotted lines demarcate assay lower limit of linear performance range (log10 of 2.65 corresponding to 450 copies/mL); positive values below this limit are plotted as 450 copies/ml. Open symbols represent animals with viral loads below the limit of detection of the assay. Box plot horizontal lines indicate the mean; top and bottom reflect the minimum and maximum. Significant differences between control and vaccinated animals at day 2 post-challenge are indicated. Significance was assessed using a Kruskal-Wallis test followed by a Dunn’s post-test.

Viral replication was detected in saliva in all control animals on day 1 and persisted in five animals through day 4 (Fig. 3*C*). Values were lower than those in BAL or NP swabs and tapered to undetectable levels more rapidly. Overall fewer vaccinated animals contained sgmRNA in their saliva and replicating virus was detected in only one animal from each vaccine dose group starting on day 2. The kinetics of SARS-CoV-2 total RNA, which is more likely to reflect material from the viral inoculum, paralleled results described above for sgmRNA in BAL, NP swabs and saliva (Fig. S4).

### Respiratory tract pathology and antigen expression

Vaccine efficacy was also assessed by histopathologic analysis of lung tissue from 3-5 macaques from each group necropsied at day 7 post-challenge. By this point, all unvaccinated animals had developed histopathologic evidence of multifocal, mild to moderate interstitial pneumonia (Fig. 4*A*). The pneumonia was characterized by type II pneumocyte hyperplasia, alveolar edema, alveolar inflammatory and necrotic debris, thickening of alveolar septae, increased numbers of pulmonary macrophages (including multinucleated giant cells), and vasculitis of small-to medium-caliber blood vessels. The middle and caudal lung lobes were most severely affected in all four unvaccinated animals. Histologic evidence of interstitial pneumonia was not observed in animals from any of the vaccinated groups (Fig. 4*B,C*). However, in each of the vaccine groups, there was minimal to mild mononuclear to mixed cellular infiltrates centered on small-to-medium caliber blood vessels. Immunohistochemistry demonstrated SARS-CoV-2 viral antigen in small numbers of alveolar pneumocytes and pulmonary macrophages in at least one lung section of every unvaccinated animal (Fig. 4*D*). No viral antigen was detected in the lungs of any of the animals in any of the vaccine groups (Fig. 4*E,F*). Overall, pathological findings were significantly reduced by vaccination (Fig. 4*G*). No significant histopathologic differences were observed between vaccinated and unvaccinated animals at day 14, consistent with transient SARS-CoV-2 pathology in this model. Mild perivascular infiltrates occasionally remained in some animals from all groups. In sum, vaccination with 5 or 50 μg RFN prevented moderate disease and viral protein expression in the lungs.

**Figure 4.**
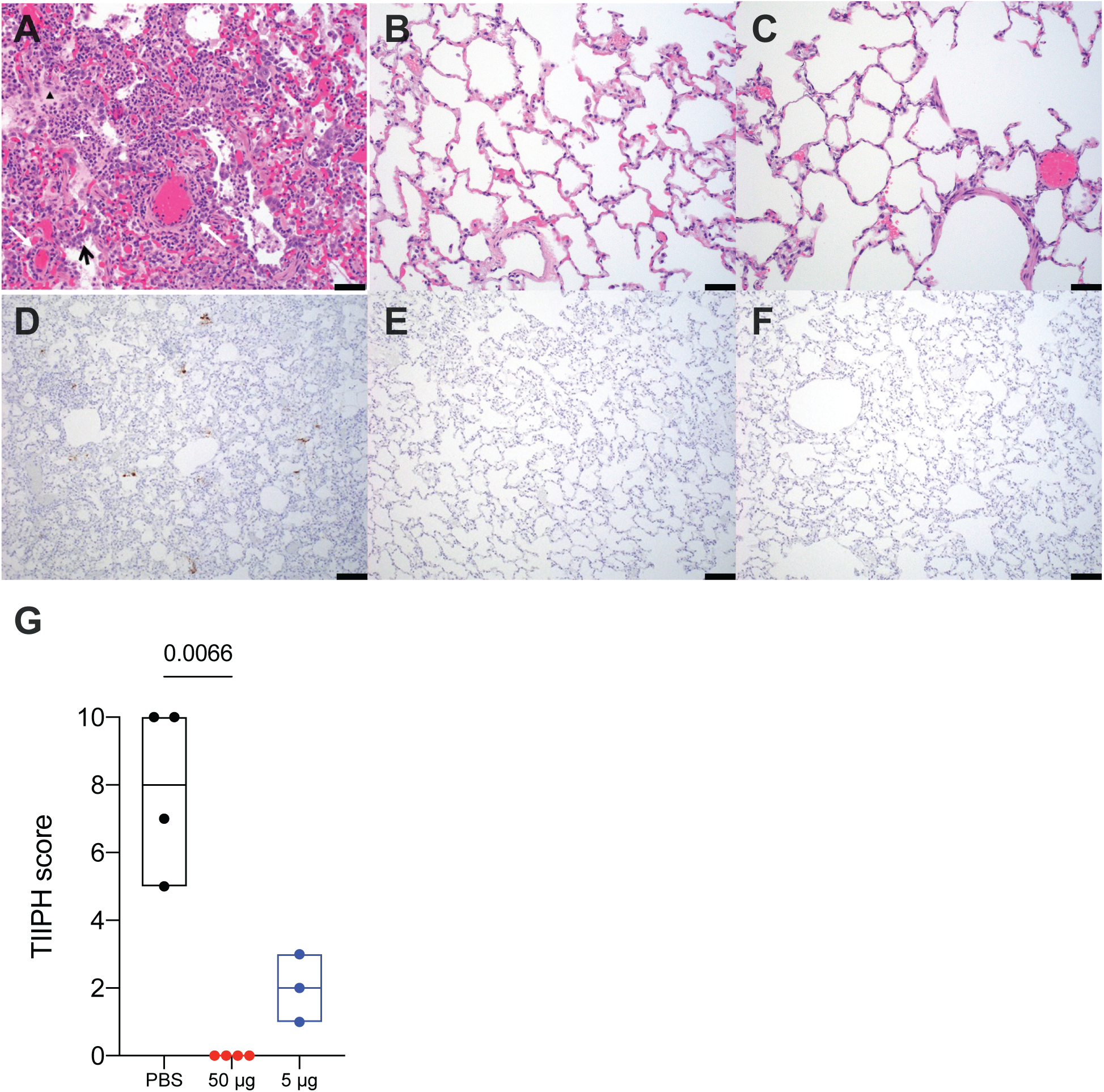
Histopathology and virus detection in the lungs following SARS-CoV-2 challenge. Lung parenchymal tissue was assessed for pathology and viral antigen seven days post-challenge. (*A-C*) Paraffin-embedded sections were stained with hematoxylin and eosin and shown for animals that received PBS *(A)*, 50 µg RFN *(B)* and 5 μg RFN *(C)*. Inflammatory debris (white star), type II pneumocyte hyperplasia (black arrow), edema (triangle) and vasculitis of small-to medium-caliber blood vessels (white arrows) is indicated. Scale bars represent 50 µm. (*D-F*) SARS-CoV-2 nucleocapsid detected by immunohistochemistry in alveolar pneumocytes, pulmonary macrophages and endothelial cells, appears as brown aggregates. Scale bars represent 100 µm. Representative images are shown. (*G*) Each pathologic finding was quantified for six lung sections and reported as a combined TIIPH score for all animals necropsied at day seven post-challenge.

### Cross-reactive immunity to emergent SARS-CoV-2 variants and SARS-CoV-1

Given concerns about circulating SARS-CoV-2 viral variants’ increased resistance to currently available vaccines, we assessed RFN vaccinated macaque serum for neutralizing antibody responses against two variants of concern, B.1.1.7 and B.1.351. In an authentic virus neutralization assay, reciprocal neutralization ID50 GMT titers against B.1.1.7 were 73,983 two weeks following the second 50 μg dose (Fig. 5*A*). This translated to ∼3.8-fold greater titers than those against the wild-type, vaccine-matched USA-WA1 strain. Titers against the two strains were similar when measured by the pseudovirus neutralization assay (Fig. 5*B*). These trends were observed regardless of vaccine dose, though responses were slightly lower with 5 µg RFN. GMT neutralization titers against B.1.351 decreased approximately 2-fold to 8,070 and 9,876 in the 50 μg dose group in the authentic virus and pseudovirus assays respectively, indicating only a minor diminution in potency compared to USA-WA1 (Fig. 5*A-C*). Thus, RFN vaccination elicited broadly reactive neutralizing antibody responses with potent activity against two important variants. Serum binding to the variant forms of SARS-CoV-2 was also assessed by biolayer interferometry (Fig. S6). In both vaccine groups no changes in binding were observed to B.1.1.7 (N501Y mutation), while responses to B.1.351 (K417N, E484K, N501Y mutations) trended ∼15% lower in both vaccine groups.

**Figure 5.**
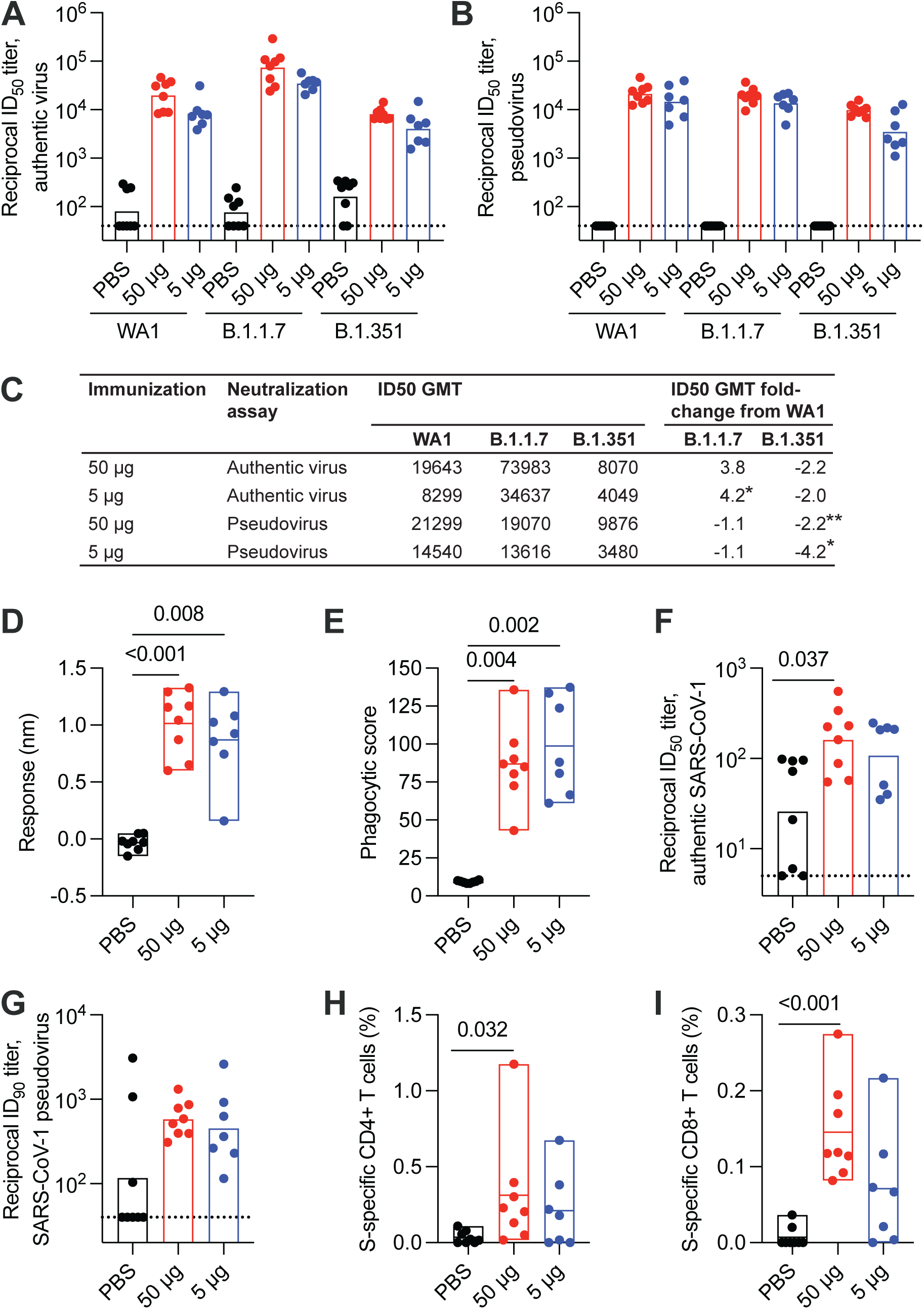
Cross-reactive immune responses to SARS-CoV-2 variants and SARS-CoV-1. Serum and PBMC collected two weeks after the last vaccination was assessed for cross-reactivity to variants of concern and SARS-CoV-1. *(A)* Authentic virus and *(B)* pseudovirus neutralizing antibody responses to variants B.1.1.7 and B.1.351. Corresponding responses to SARS-CoV-2 USA-WA1 authentic virus and Wuhan-1 pseudovirus are shown. Bars indicate the geometric mean titer. *(C)* Reciprocal ID50 geometric mean titer (GMT) fold-change from wild-type neutralization (WA1 or Wuhan-1) with statistical significance set at a p-value of < 0.05. *(D)* Serum binding responses to SARS-CoV-1 RBD assessed by biolayer interferometry. *(E)* Antibody-dependent cellular phagocytosis of SARS-CoV-1 spike trimer-coated fluorescent beads. *(F)* Authentic SARS-CoV-1 (Urbani) neutralization titers (ID50). *(G)* SARS-CoV-1 (Urbani) pseudovirus neutralization (ID90). *(H)* SARS-CoV-1 (Urbani) S-specific memory CD4+ Th1 and *(I)* CD8+ responses assessed by peptide pool stimulation and ICS (INF*γ*, IL-2 and TNF). Significance was assessed with a Kruskal-Wallis test followed by a Dunn’s post-test. Bars indicate the geometric mean titer.

In addition to SARS-CoV-2 variants of concern, another open question in the field is the ability of existing SARS-CoV-2 vaccine platforms to protect against future SARS-like coronavirus outbreaks. Cross-protective vaccine-elicited immunity against SARS-CoV-1 may be a useful metric to address this question. We measured IgG antibody responses able to bind SARS-CoV-1 RBD by biolayer interferometry in macaque serum at week 2 following the second vaccination. All RFN vaccinated animals developed cross-reactive binding antibodies to SARS-CoV-1 at levels approximately half those to SARS-CoV-2 (Fig. 5*D*, Fig. S1). Binding responses were also measured to a series of SARS-CoV-1 and SARS-CoV-2 antigens using a Luminex assay (Fig. S5), with strong binding responses observed to the SARS-1 S1 subunit and RBD, but not against the S2 subunit or N-terminal domain. SARS-CoV-1 RBD-specific binding antibody responses were ∼70% that of the SARS-CoV-2 response. The functional capacity of these cross-binding antibodies to mediate effector activity was assessed in an ADCP assay using SARS-CoV-1 trimeric S antigen. SARS-CoV-1 ADCP responses were observed in plasma of all vaccinated animals and were comparable between the dose groups (Fig. 5*E*).

Neutralizing titers against SARS-CoV-1 were measured using both authentic virus and pseudovirus neutralization assays, with cross-neutralizing responses observed in most RFN vaccinated animals (Fig. 5*F,G*). Significant authentic virus neutralization titers were elicited by 50 μg RFN two weeks following the second immunization. SARS-CoV-1 pseudovirus neutralization activity was also observed in both the 50 and 5 μg groups, though background in a subset of control animals limited interpretation of both assays.

To assess T cell immunity cross-reactivity to SARS-CoV-1, we evaluated whether the RFN vaccine-elicited T cells could recognize SARS-CoV-1 S. PBMC stimulated with SARS-CoV-1 S peptide pools were stained for intracellular cytokine expression to quantitate cross-reactive T cells. Significant CD4+ T cell Th1 responses were observed following the 50 μg RFN vaccination series, though they were ∼5-fold lower in magnitude than those to SARS-CoV-2 S (Fig. 5*H*). SARS-CoV-1 S-specific CD40L responses were comparable to the Th1 responses for both dose groups (Fig. S7*A*). IL-21 and Th2 CD4+ T cell responses were minimal or negligible (Fig. S7*B,C*). Significant cross-reactive CD8+ T cells were elicited and similar in magnitude to SARS-CoV-2-specific responses (∼0.1-0.3%) (Fig. 5*I*), suggesting that the CD8+ T cell RBD epitope specificities elicited by RFN vaccination may be relatively conserved. Again, responses trended greater with the higher vaccine dose. These data indicate that S-specific CD4+ and CD8+ T cells generated by ALFQ-adjuvanted RFN were able to cross-react with sequence divergent SARS-CoV-1.

## DISCUSSION

New SARS-CoV-2 vaccines may be needed to address concerns regarding emerging virus variants less sensitive to immunity elicited by current vaccines (1, 44–47). In this study, we evaluated a candidate RFN vaccine adjuvanted with ALFQ in rhesus macaques and observed robust and broad humoral and T cell responses and protection from high-dose respiratory tract challenge. Binding, neutralizing, and effector antibody responses were elicited in all animals and of comparable or greater magnitude to that observed in pre-clinical studies of EUA vaccines (15, 48, 49). Spike-specific CD4+ T cell responses exceeded 0.5% of memory cells and were predominantly of the Th1 phenotype. Using a rigorous challenge model in which viral loads of control animals exceeded that of published macaque studies, replicating virus was rapidly cleared in the airways of vaccinated animals. Cross-reactive antibody responses were either higher or similar against the B.1.1.7 VOC in authentic and pseudovirus neutralization assays, while B.1.351 reactivity was diminished by only ∼2-fold. Additionally, binding and functional antibodies were also reactive to SARS-CoV-1, which is 36% amino acid sequence divergent from SARS-CoV-2 in the RBD (50). Overall, these data indicate broad, potent and efficacious immunity elicited by RFN-ALFQ.

This study provides strong evidence that RBD-directed vaccination in primates is able to protect against SARS-CoV-2 infection and elicit neutralization breadth against variant B.1.351, which has shown the greatest resistance to neutralization by vaccinee sera (19, 51, 52). While many RBD-based immunogens have been shown to be immunogenic in small and large animal models (24–27), limited studies assessed efficacy against viral challenge and neutralization activity against VOC. A recent macaque study investigated the immunogenicity and protective efficacy of a three-dose regimen of an RBD-ferritin nanoparticle-based vaccine (RBD scNP) and reported efficacy upon challenge, with no sgmRNA detected in the upper or lower airways of vaccinated animals at day 2 post-challenge (53). Neutralizing antibody titers were observed for the B.1.1.7 variant but were not assessed for B.1.351. Here, RFN vaccination elicited similar B.1.1.7 neutralization as RBD scNP, but also neutralized B.1.351 with minimal reduction in potency relative to USA-WA1. In another study, vaccination with RBD fused to the Fc domain of human IgG1 reduced total viral RNA following challenge in cynomolgus macaques, though virus replication was not assessed (54). Our findings demonstrate RBD-specific immunity elicited by a condensed two-dose vaccine regimen is protective and, importantly, cross-neutralizes the more resistant B.1.351 variant.

The immunogenicity and efficacy of ALFQ-adjuvanted RFN compares favorably to pre-clinical macaque data reported for three COVID-19 vaccines authorized for emergency use. RFN vaccination elicited peak mean SARS-CoV-2 pseudovirus neutralization reciprocal titers of 14,540 and 21,298 for the 5 and 50 µg groups, respectively, exceeding those of the EUA vaccines, which ranged from 408-1,862 (48, 49, 55). While neutralizing activity is unlikely to be the sole determinant of vaccine-mediated protection, it has been predictive of efficacy in human trials (1). Therefore, ∼10-50-fold greater neutralizing titers by RFN relative to those elicited in NHP studies by efficacious vaccines currently in clinical use strongly suggests that RFN would be protective in humans. In addition, breadth against the B.1.351 VOC appears greater than that of EUA vaccines in humans, as the modest ∼2-fold reduction in B.1.351 neutralization activity relative to wild-type virus reported here is less than the ∼6-12-fold reduction in mRNA vaccinee sera (19). The most advanced platform closest in design and composition to RFN is NVX-CoV2373, a prefusion spike nanoparticle vaccine delivered with a saponin-based Matrix-M adjuvant. NVX-CoV2373 elicited neutralizing antibody titers of 6,400-17,000 in macaques (56, 57), greater than those achieved by current EUA vaccines and less than or similar to those elicited by RFN. T cell immunity was also more pronounced with RFN. S-specific Th1 CD4+ T cells ranged from 0.5-5% following 50 µg RFN, compared to peak values of 0.1-0.2% in NHP vaccinated with the EUA vaccines (48, 49, 55).

When considering vaccination strategies for clinical use in humans, cost and manufacturability are important considerations. The comparison of low or high dose vaccine regimens presented here demonstrated that immune responses did not significantly differ between the 5 and 50 µg doses, although the 50 μg group did trend towards higher responses and a slightly earlier resolution of viral load. The power to detect these differences may have been limited by small sample sizes. It is likely that as doses decrease further protective efficacy will wane, and such experiments may allow further elucidation of correlates of protection. This absence of a strong dose titration effect suggests that vaccination with the lower dose may be possible for dose sparing purposes, although clinical testing and assessment of response durability are required.

In addition to the RFN vaccine described here, we have also developed a similar ferritin nanoparticle immunogen displaying the full prefusion stabilized SARS-CoV-2 spike glycoprotein (SpFN) and reported its immunogenicity and efficacy in non-human primates (58). Compared to two-dose SpFN regimens, RFN elicited binding and neutralizing antibody and T cell responses of a similar magnitude, albeit with a trend towards slightly lower titers. Post-challenge control of viral replication was also similar, though viral clearance after SpFN vaccination trended faster from the BAL at day 2 post-challenge and from NP swabs by day 4. The overall magnitude of these differences was small and suggests that both the RBD and S proteins are similarly immunogenic and protective when complexed to ferritin nanoparticles and administered with ALFQ adjuvant at these vaccine doses. However, S-based immunogens may offer the advantage of broadening the specificity of the immune response to other domains and subdomains of the spike protein, limiting potential for viral escape. These findings support further clinical development of both products.

There exists a strong potential for future pandemics arising from zoonotic SARS-like betacoronaviruses entering into humans. We report SARS-CoV-2 RFN vaccine-elicited responses that cross-react with the S glycoprotein SARS-CoV-1, including binding antibody titers within an order of magnitude of those to SARS-CoV-2. The observed cross-neutralizing and binding reactivity to SARS-CoV-1 suggests that adjuvanted RFN may be a viable candidate for vaccination against future betacoronavirus outbreaks. Work is ongoing to elucidate the potential mechanisms of cross-protective responses in this study, including epitope mapping of the antibody binding responses. Taken together, these findings support continued development of RFN vaccines for managing COVID-19 and SARS-like betacoronavirus outbreaks.

## MATERIALS AND METHODS

### Vaccine and adjuvant production

#### DNA plasmid construction and preparation

The SARS-CoV-2 RBD-ferritin construct was derived from the Wuhan-Hu-1 strain genome sequence (GenBank MN9089473) comprising residues 331 – 527. RBD was attached to *Helicobacter pylori* ferritin using a GSGGGG linker followed by a short sequence (ESQVRQQFSK) derived from bullfrog ferritin (59) and synthesized by GenScript, to include a N-terminal hexa-histadine (his) tag for purification. Additional point mutations (Y453R, L518N, L519K, H520S) were introduced in the RBD, using a modified QuikChange site-directed mutagenesis protocol (Agilent Technologies, Santa Clara, CA) and designated as construct RFN_131. The construct used a prolactin leader (PL) sequence (60). Plasmid DNA generated by site-directed mutagenesis was prepared from *Eschericia coli* Stbl3 cells. Large-scale DNA isolation was performed using either endo free Maxiprep, Megaprep or Gigaprep kits (Qiagen, Hilden, Germany).

#### Immunogen expression and purification

SARS-CoV-2 RFN_131 immunogen (RFN) was expressed in 293Expi mammalian cell lines by transient transfection using Turbo293 transfection reagent (Speed Biosystems, Gaithersburg, MD). Expression cultures were grown in polycarbonate baffled shaker flasks at 34°C and 8% CO2 at 120 rpm. Cells were harvested five days post-transfection via centrifugation at 3,500 x g for 30 min. Culture supernatants were filtered with a 0.22-µm filter and stored at 4°C prior to purification. RFN was purified using Ni-NTA affinity chromatography. 1 mL Ni-NTA resin (Thermo Scientific) was used to purify protein from 1 L of expression supernatant. Ni-NTA resin was equilibrated with 5 column volumes (CV) of phosphate buffered saline (PBS) (pH 7.4) followed by supernatant loading at room temperature. Unbound protein was removed by washing with 200 CV of PBS, followed by 50 CV 10mM imidazole in PBS. Bound protein was eluted with 220mM imidazole in PBS. Purification purity was assessed by SDS-PAGE. RFN was concentrated in the presence of 5% glycerol and then further purified by size-exclusion chromatography using a 16/60 Superdex-200 purification column. Endotoxin levels for ferritin nanoparticle immunogens were evaluated (Endosafe nexgen-PTS, Charles River Laboratories) and 5 % v/v glycerol was added prior to filter-sterilization with a 0.22-µm filter, flash-freezing in liquid nitrogen, and storage at −80°C. Ferritin nanoparticle formation was confirmed by negative-stain electron microscopy and dynamic light scattering by determining the hydrodynamic diameter at 25 °C using a Malvern Zetasizer Nano S (Malvern, Worcestershire, UK) equipped with a 633-nm laser.

#### Adjuvant preparation

Army Liposomal Formulation with QS21 (ALFQ), formulation was prepared as previously described (61, 62). ALFQ is a unilamellar liposome comprised of dimyristoyl phosphatidylcholine (DMPC), dimyristoyl phosphatidylglycerol (DMPG), cholesterol (Chol), and synthetic monophosphoryl lipid A (3D-PHAD^®^) (Avanti Polar Lipids, Alabaster, AL) and QS-21 (Desert King, San Diego, CA). DMPC and cholesterol were dissolved in chloroform and DMPG and 3D-PHAD^®^ were dissolved in chloroform:methanol 9:1. Lipids were mixed in a molar ratio of 9:1:12.2:0.114 (DMPC:DMPG:Chol:3D-PHAD^®^) and the solvent was removed by rotary evaporation. Lipids were suspended in Sorenson’s PBS, pH 6.2, microfluidized to form small unilamellar vesicles and filtered. QS-21 was solubilized in Sorenson’s PBS, pH 6.2, filtered and added to the vesicles to form ALFQ. The final lipid ratio was 9:1:12.2:0.114:0.044 (DMPC:DMPG:Chol:3D-PHAD^®^:QS-21).

#### Immunogen formulation

RFN was diluted in dPBS to 0.1 mg/mL or 0.01 mg/mL and mixed 1:1 with 2X ALFQ on a tilted slow-speed roller at room temperature for 10 min, followed by incubation at 4°C for 50 min. Reagents were equilibrated to room temperature before use and immunizations were performed within 4 h of vaccine formulation. Each vaccine comprised a 1.0 mL solution of RFN formulated with ALFQ. The 3D-PHAD^®^ and QS-21 doses were 200 and 100 µg, respectively.

### Study design and procedures

Twenty-three male and female specific-pathogen-free, research-naïve Chinese-origin rhesus macaques (age 3 - 7 years) were distributed—on the basis of age, weight and sex—into 3 cohorts of 7-8 animals (Table S1). Animals were vaccinated intramuscularly with either 50 or 5 μg of RFN, formulated with ALFQ, and control group animals received 1 mL of PBS, in the anterior proximal quadriceps muscle, on alternating sides with each dose in the series. Immunizations were administered twice 4 weeks apart. Animals were challenged 4 weeks after the second immunization with virus stock obtained through BEI Resources, NIAID, NIH: SARS-Related Coronavirus 2, Isolate USA-WA1/2020, NR-53780 (Lot# 70038893). Virus was stored at −80°C prior to use, thawed by hand and placed immediately on wet ice. Stock was diluted to 5×10^5^ TCID50/mL in PBS and vortexed gently for 5 sec prior to inoculation via combined intratracheal and intranasal routes (1 mL each).

All procedures were carried out in accordance with institutional, local, state and national guidelines and laws governing animal research included in the Animal Welfare Act. Animal protocols and procedures were reviewed and approved by the Animal Care and Use Committee of both the US Army Medical Research and Development Command (USAMRDC, protocol 11355007.03) Animal Care and Use Review Office as well as the Institutional Animal Care and Use Committee of Bioqual, Inc. (protocol number 20-092), where nonhuman primates were housed for the duration of the study. USAMRDC and Bioqual, Inc. are both accredited by the Association for Assessment and Accreditation of Laboratory Animal Care and are in compliance with the Animal Welfare Act and Public Health Service Policy on Humane Care and Use of Laboratory Animals.

### Convalescent Plasma Samples

A panel of 41 human convalescent-phase plasma samples were obtained from BEI Resources Repository (N=30), StemExpress (East Norritin, PA) (N=7) and a Walter Reed Army Institute of Research institutional review board-approved leukapheresis protocol (#1386H) (N=4) for which written informed consent was provided by participants. Samples were collected from males (N=20) and females (N=21) ranging in age from 31 to 71 years. Individuals donated plasma specimens approximately four-to-eight weeks after laboratory-confirmed SARS-CoV-2 infection and had histories of asymptomatic-to-mild-to-moderate clinical presentation.

### Antibody responses

#### Binding antibodies

SARS-CoV-2-specific binding IgG antibodies and ACE2 inhibiting antibodies were measured using MULTI-SPOT^®^ 96-well plates, (Meso Scale Discovery (MSD), Rockville, MD). Multiplex wells were coated with SARS-CoV-2 antigens, S and RBD, at a concentration of 200-400 ng/mL and BSA, which served as a negative control. Four-plex MULTISPOT plates were blocked with MSD Blocker A buffer for 1 h at room temperature (RT) while shaking at 700 rpm. Plates were washed with buffer before the addition of reference standard and calibrator controls. Serum samples were diluted at 1:1,000 - 1:100,000 in diluent buffer, then added to each of four wells. Plates were incubated for 2 h at room temperature while shaking at 700 rpm, then washed. MSD SULFO-TAG^TM^ anti-IgG antibody was added to each well. Plates were incubated for 1 h at RT with shaking at 700 rpm and washed, then MSD GOLD^TM^ Read buffer B was added to each well. Plates were read by the MESO SECTOR SQ 120 Reader. IgG concentration was calculated using DISCOVERY WORKBENCH^®^ MSD Software and reported as arbitrary units (AU)/mL.

For binding antibodies that block S or RBD binding to ACE2, antigen-coated plates were blocked and washed as described above. Assay calibrator and samples were diluted at 1:25 - 1:1,000 in MSD Diluent buffer, then added to the wells. Plates were incubated for 1 h at room temperature while shaking at 700 rpm. ACE2 protein conjugated with MSD SULFO-TAG^TM^ was added and plates were incubated for 1 h at room temperature while shaking at 700rpm and washed and read as described above.

Binding antibody measurements by octet biolayer interferometry were made using HIS1K biosensors hydrated in PBS prior to use, using an Octet FortéBio Red96 instrument (Sartorius, Fremont CA). All assay steps were performed at 30°C with agitation set at 1,000 rpm. Baseline equilibration of the HIS1K biosensors (Sartorius, Fremont, CA)) was carried out with PBS for 15 sec, prior to SARS-CoV2 RBD molecules (30 µg/mL diluted in PBS) loading for 120 sec. Biosensors were dipped in assay buffer (15 sec in PBS), dipped in the serum samples (100-fold dilution) for 180 sec, and binding response (nm) was recorded for 180 sec.

### Virus neutralization

#### SARS-CoV-1 and SARS-CoV-2 pseudovirus neutralization

The S expression plasmid sequence for SARS-CoV-2 was codon optimized and modified to remove an 18 amino acid endoplasmic reticulum retention signal in the cytoplasmic tail to improve S incorporation into pseudovirions (PSV). PSV were produced by co-transfection of HEK293T/17 cells with a SARS-CoV-2 S plasmid (pcDNA3.4), derived from the Wuhan-Hu-1 genome sequence (GenBank accession number: MN908947.3) and an HIV-1 NL4-3 luciferase reporter plasmid (pNL4-3.Luc.R-E-, NIH HIV Reagent Program, Catalog number 3418). Infectivity and neutralization titers were determined using ACE2-expressing HEK293 target cells (Integral Molecular, Philadelphia, PA) in a semi-automated assay format using robotic liquid handling (Biomek NXp Beckman Coulter, Brea, CA). Virions pseudotyped with the vesicular stomatitis virus (VSV) G protein were used as a non-specific control. Test sera were diluted 1:40 in growth medium and serially diluted; then 25 μL/well was added to a white 96-well plate. An equal volume of diluted PSV was added to each well and plates were incubated for 1 h at 37°C. Target cells were added to each well (40,000 cells/well) and plates were incubated for an additional 48 h. Relative light units (RLU) were measured with the EnVision Multimode Plate Reader (Perkin Elmer, Waltham, MA) using the Bright-Glo Luciferase Assay System (Promega, Madison, WI). Neutralization dose–response curves were fitted by nonlinear regression using the LabKey Server. Final titers are reported as the reciprocal of the serum dilution necessary to achieve 50% inhibition SARS-CoV-2 (ID50, 50% inhibitory dose) or 90% inhibition for SARS-CoV-1 (ID90, 90% inhibitory dose). Assay equivalency was established by participation in the SARS-CoV-2 Neutralizing Assay Concordance Survey run by the Virology Quality Assurance Program and External Quality Assurance Program Oversite Laboratory at the Duke Human Vaccine Institute.

#### Authentic SARS-CoV-2 wild-type neutralization assay

Authentic virus neutralization was measured using SARS-CoV-2 (2019-nCoV/USA_WA1/2020) obtained from the Centers for Disease Control and Prevention and passaged once in Vero CCL81 cells (ATCC). Rhesus sera were serially diluted and incubated with 100 focus-forming units of SARS-CoV-2 for 1 h at 37°C. Serum-virus mixtures were added to Vero E6 cells in 96-well plates and incubated for 1 h at 37°C. Cells were overlaid with 1% (w/v) methylcellulose in MEM. After 30 h, cells were fixed with 4% PFA in PBS for 20 min at room temperature then washed and stained overnight at 4°C with 1 µg/mL of CR3022 antibody in PBS supplemented with 0.1% saponin and 0.1% bovine serum albumin. Cells were subsequently stained with HRP-conjugated goat anti-human IgG for 2 h at room temperature. SARS-CoV-2-infected cell foci were visualized with TrueBlue peroxidase substrate (KPL) and quantified using ImmunoSpot microanalyzer (Cellular Technologies). Neutralization curves were generated with Prism software (GraphPad Prism 8.0).

#### Authentic SARS-CoV-2 variant and SARS-CoV-1 neutralization assay

The SARS-CoV-2 viruses USA-WA1/2020 (WA1), USA/CA_CDC_5574/2020 (B.1.1.7), and hCoV-19/South Africa/KRISP-EC-K005321/2020 (B.1.351) were obtained from BEI Resources (NIAID, NIH) and propagated for one passage using Vero-E6 cells. Virus infectious titer was determined by an end-point dilution and cytopathic effect (CPE) assay on Vero-E6 cells. In brief, serum samples were heat inactivated and subjected to successive 3-fold dilutions starting from 1:50. Triplicates of each dilution were incubated with SARS-CoV-2 at an MOI of 0.1 in EMEM with 7.5% inactivated fetal calf serum (FCS) for 1 h at 37°C. Virus-antibody mixture was transferred onto a monolayer of Vero-E6 cells grown overnight and incubated for ∼70 h. CPE of viral infection was visually scored for each well in a blinded fashion by two independent observers. Results were reported as percentage of neutralization at a given sample dilution. A SARS-CoV-1 authentic plaque reduction virus neutralization assay was performed similarly to previously described (63) with the following modifications. The starting dilution of serum was 1:5 and ∼100 PFU of virus was used for virus/serum incubation. The overlay used after virus adsorption was DMEM containing 2% FBS and 20% methylcellulose. Plates were then incubated for 5 days, and post crystal violet staining the washing step utilized water. Plaques were graded as follow: ∼50 plaques/50% MD (+); ∼75 plaques/75% MD (++); ∼100 plaques/100% MD (+++); ∼25 plaques/25%. All negative control wells were solid monolayers.

#### Antibody-dependent neutrophil phagocytosis (ADNP)

Biotinylated SARS-CoV-2 prefusion stabilized S trimer was incubated with yellow-green streptavidin-fluorescent beads (Molecular Probes, Eugene, OR) for 2 h at 37°C. 10 μL of a 100-fold dilution of protein-coated beads was incubated for 2 h at 37°C with 100 μL of 8,100-fold diluted plasma samples before addition of effector cells (50,000 cells/well). Fresh human peripheral blood mononuclear cells were used as effector cells after red blood cell lysis with ACK lysing buffer (ThermoFisher Scientific, Waltham, MA). After 1 h incubation at 37°C, cells were washed, surface stained, fixed with 4% formaldehyde solution (Tousimis, Rockville, MD) and fluorescence was evaluated on an LSRII flow cytometer (BD Bioscience, San Jose, CA). Antibodies used for flow cytometry included anti-human CD3 AF700 (clone UCHT1), anti-human CD14 APC-Cy7 (clone MϕP9) (BD Bioscience, San Jose, CA) and anti-human CD66b Pacific Blue (clone G10F5) (Biolegend, San Diego, CA). Phagocytic score was calculated by multiplying the percentage of bead-positive neutrophils (SSC high, CD3-CD14-CD66+) by the geometric mean of the fluorescence intensity of bead-positive cells; and dividing by 10,000.

#### Antibody-dependent cellular phagocytosis (ADCP)

ADCP was measured as previously described (64). Briefly, biotinylated SARS-CoV-1 or SARS-CoV-2 prefusion-stabilized S trimer was incubated with red streptavidin-fluorescent beads (Molecular Probes, Eugene, OR) for 2 h at 37°C. 10 μL of a 100-fold dilution of beads–protein was incubated for 2 h at 37°C with 100 μL of 8,100-fold (SARS-CoV-2) or 900-fold (SARS-CoV-1) diluted plasma samples before addition of THP-1 cells (20,000 cells per well; Millipore Sigma, Burlington, MA). After 19 h incubation at 37°C, the cells were fixed with 2% formaldehyde solution (Tousimis, Rockville MD) and fluorescence was evaluated on an LSRII flow cytometer (BD Bioscience, San Jose, CA). The phagocytic score was calculated by multiplying the percentage of bead-positive cells by the geometric mean of the fluorescence intensity of bead-positive cells and dividing by 10,000.

#### Opsonization

SARS-CoV-2 Spike-expressing expi293F cells were generated by transfection with linearized plasmid (pcDNA3.1) encoding codon-optimized full-length SARS-CoV-2 Spike protein matching the amino acid sequence of the IL-CDC-IL1/2020 isolate (GenBank ACC# MN988713). Stable transfectants were single-cell sorted and selected to obtain a high-level Spike surface expressing clone (293F-Spike-S2A). SARS-CoV-2 S expressing cells were incubated with 200-fold diluted plasma samples for 30 min at 37°C. Cells were washed twice and stained with anti-human IgG PE, anti-human IgM Alexa Fluor 647, and anti-human IgA FITC (Southern Biotech, Birmingham, AL). Cells were then fixed with 4% formaldehyde solution and fluorescence was evaluated on an LSRII flow cytometer (BD Bioscience, San Jose, CA).

#### Antibody-dependent complement deposition (ADCD)

ADCD was adapted from methods described previously (65). Briefly, SARS-CoV-2 S expressing expi293F cells were incubated with 10-fold diluted, heat-inactivated (56°C for 30 min) plasma samples for 30 min at 37°C. Cells were washed twice and resuspended in R10 media. During this time, lyophilized guinea pig complement (CL4051, Cedarlane, Burlington, Canada) was reconstituted in 1 mL cold water and centrifuged for 5 min at 4°C to remove aggregates. Cells were washed with PBS and resuspended in 200 μL of guinea pig complement, which was prepared at a 1:50 dilution in Gelatin Veronal Buffer with Ca^2+^ and Mg^2+^ (IBB-300x, Boston BioProducts, Ashland, MA). After incubation at 37°C for 20 min, cells were washed in PBS 15mM EDTA (ThermoFisher Scientific, Waltham, MA) and stained with an anti-Guinea Pig Complement C3 FITC (polyclonal, ThermoFisher Scientific, Waltham, MA). Cells were then fixed with 4% formaldehyde solution and fluorescence was evaluated on a LSRII flow cytometer (BD Bioscience, San Jose, CA).

#### Trogocytosis

Trogocytosis was measured using a previously described assay (38). Briefly, SARS-CoV-2 Spike expressing expi293F cells were stained with PKH26 (Sigma-Aldrich, St-Louis, MO). Cells were then washed with and resuspended in R10 media. Cells were then incubated with 200-fold diluted plasma samples for 30 min at 37°C. Effector peripheral blood mononuclear cells were next added to the R10 media at an effector to target (E:T) cell ratio of 50:1 and then incubated for 5 h at 37°C. After the incubation, cells were washed, stained with live/dead aqua fixable cell stain (Life Technologies, Eugene, OR) and CD14 APC-Cy7 (clone MϕP9) for 15 min at room temperature, washed again, and fixed with 4% formaldehyde (Tousimis, Rockville, MD) for 15 min at room temperature. Fluorescence was evaluated on a LSRII flow cytometer (BD Bioscience, San Jose, CA). Trogocytosis was evaluated by measuring the PKH26 mean fluorescence intensity of the live CD14+ cells.

### Antigen-specific T cell responses

Cryopreserved peripheral blood mononuclear cells were thawed and rested for 6 h in R10 with 50 U/mL Benzonase Nuclease (Sigma-Aldrich, St. Louis, MO). They were then stimulated with peptide pools for 12 h. Stimulations consisted of two pools of peptides spanning the Spike protein of SARS-CoV-2 or SARS-CoV-1 (1 µg/mL, JPT, PM-WCPV-S and PM-CVHSA-S respectively) in the presence of Brefeldin A (0.65 µL/mL, GolgiPlug^TM^, BD Cytofix/Cytoperm Kit, Cat. 555028), co-stimulatory antibodies anti-CD28 (BD Biosciences Cat. 555725 1 µg/mL) and anti-CD49d (BD Biosciences Cat. 555501; 1ug/mL) and CD107a (H4A3, BD Biosciences Cat. 561348, Lot 9143920 and 253441). Following stimulation, cells were stained serially with LIVE/DEAD Fixable Blue Dead Cell Stain (ThermoFisher #L23105) and a cocktail of fluorescent-labeled antibodies (BD Biosciences unless otherwise indicated) to cell surface markers CD4-PE-Cy5.5 (S3.5, ThermoFisher #MHCD0418, Lot 2118390 and 2247858), CD8-BV570 (RPA-T8, BioLegend #301038, Lot B281322), CD45RA BUV395 (5H9, #552888, Lot 154382 and 259854), CD28 BUV737 (CD28.2, #612815, Lot 0113886), CCR7-BV650 (GO43H7, # 353234, Lot B297645 and B316676) and HLA-DR-BV480 (G46-6, # 566113, Lot 0055314). Intracellular cytokine staining was performed following fixation and permeabilization (BD Cytofix/Cytoperm, BD Biosciences) with CD3-Cy7APC (SP34-2, #557757, Lot 6140803 and 121752), CD154-Cy7PE (24-31, BioLegend # 310842, Lot B264810 and B313191), IFN*γ*-AF700 (B27, # 506516, Lot B187646 and B290145), TNF*α*-FITC (MAb11, # 554512, Lot 15360), IL-2-BV750 (MQ1-17H12, BioLegend #566361, Lot 0042313), IL-4 BB700 (MP4-25D2, Lot 0133487 and 0308726), MIP-1b (D21-1351, # 550078, Lot 9298609), CD69-ECD (TP1.55.3, Beckman Coulter # 6607110, Lot 7620070 and 7620076), IL-21-AF647 (3A3-N2.1, # 560493, Lot 9199272 and 225901), IL-13-BV421 (JES10-5A2, # 563580, Lot 9322765, 210147 and 169570) and IL-17a-BV605 (BL168, Biolegend #512326, B289357). Sample staining was measured on a FACSymphony™ A5 SORP (Becton Dickenson) and data was analyzed using FlowJo v.9.9 software (Tree Star, Inc.). CD4+ and CD8+ T cell subsets were pre-gated on memory markers prior to assessing cytokine expression as follows: single-positive or double-negative for CD45RA and CD28. Boolean combinations of cells expressing one or more cytokines were used to assess the total S-specific response of memory CD4+ or CD8+ T cells. Responses from the two-peptide pools spanning SARS-CoV-2 S or SARS-CoV-1 S were summed. Display of multicomponent distributions were performed with SPICE v6.0 (NIH, Bethesda, MD).

### Total and subgenomic messenger (sgm) RNA quantification

Real-time quantitative PCR was carried out for subgenomic messenger RNA (sgmRNA) and viral load RNA quantification from NP swab, BAL fluid and saliva samples. Primers targeted the envelope (E) gene of SARS-CoV-2 (Table S2). RNA was extracted from 200 μL of specimen using the EZ1 DSP Virus kit (Qiagen) on the EZ1 Advanced XL instrument (Qiagen). Briefly, samples were lysed in 200 μL of ATL buffer (Qiagen) and transferred to the Qiagen EZ1 for extraction. Bacteriophage MS2 (ATCC, Manassas, VA) was added to the RNA carrier and used as an extraction control to monitor efficiency of RNA extraction and amplification (66). Purified RNA was eluted in 90 μL elution buffer (AVE). The RT-qPCR amplification reactions were performed in separate wells of a 96-well Fast plate for the 3 targets: sgmRNA, RNA viral load, and MS2 RNA using 10 μL of extracted material 0.72 μM of primers, 0.2 μM of probe and 1x TaqPath^TM^ 1-Step RT-qPCR (Life Technologies, Thermo Fisher Scientific, Inc.). Amplification cycling conditions were: 2 min at 25°C, 15 min at 50°C, 2 min at 95°C and 45 cycles of 3 sec at 94°C and 30 sec at 55°C with fluorescence read at 55°C. An RNA transcript for the SARS-CoV-2 E gene was used as a calibration standard. RNA copy values were extrapolated from the standard curve and multiplied by 45 to obtain RNA copies/mL. A negative control (PBS) and two positive controls, contrived using heat-inactivated SARS-CoV-2 (ATCC, VR-1986HK), at 10^6^ and 10^3^ copies/mL, were extracted and used to assess performance of both assays.

### Histopathology

Formalin-fixed sections of lung tissue were evaluated by light microscopy and immunohistochemistry. Lungs were perfused with 10% neutral-buffered formalin. Lung sections were processed routinely into paraffin wax, then sectioned at 5 µm, and resulting slides were stained with hematoxylin and eosin. Immunohistochemistry (IHC) was performed using the Dako Envision system (Dako Agilent Pathology Solutions, Carpinteria, CA, USA). Briefly, after deparaffinization, peroxidase blocking, and antigen retrieval, sections were covered with a mouse monoclonal anti-SARS-CoV nucleocapsid protein (#40143-MM05, Sino Biological, Chesterbrook, PA, USA) at a dilution of 1:4,000 and incubated at room temperature for 45 min. They were rinsed, and the peroxidase-labeled polymer (secondary antibody) was applied for 30 min. Slides were rinsed and a brown chromogenic substrate 3,3’ Diaminobenzidine (DAB) solution (Dako Agilent Pathology Solutions) was applied for 8 min. The substrate-chromogen solution was rinsed off the slides, and slides were counterstained with hematoxylin and rinsed. The sections were dehydrated, cleared with Xyless, and then cover slipped. Tissue section slides were evaluated by a board-certified veterinary anatomic pathologist who was blinded to study group allocations. Immunohistochemistry (IHC) was performed with Dako Envision. Three tissue sections from each of the right and left lung lobes were used to evaluate the lung pathology. The histopathology of each section was evaluated on a scale of 0-5: 0 - absent, 1 - minimal (<10% of tissue section affected); 2 - mild (11-25% of tissue section affected); 3 - moderate (26-50% of tissue section affected); 4 - marked (51-75% affected); 5- severe (>75% of tissue section affected). Sections were evaluated for edema, hyaline membranes, cellular infiltrates, alveolar histiocytes, type II pneumocyte hyperplasia, interstitial fibroplasia, BALT hyperplasia, bronchiolar degeneration, megakaryocytes in capillaries, and thrombosis. The scores for all six sections of each pathologic finding were combined to create the final score (TIIPH score) for individual animals.

### Statistical analysis

Primary immunogenicity outputs of binding and neutralizing antibody titers as well as T cell responses were compared across vaccination groups using the Kruskal-Wallis test. Non-parametric pairwise comparisons between groups were made using the post-hoc Dunn’s test. Statistical significance was preset at an alpha level of 0.05.

## Data Availability

All data are available in the manuscript or the supplementary materials.

## Acknowledgments

We thank M. Taddese, J. Lay, E. Zografos, J. Lynch, L. Mendez-Rivera, N. Jackson, J. Headley, M. Amare, B. Silke, U. Tran, P.J. Lee, S. Padilla, H. Hernandez, D. Coleman, H. Groove, and R.J. O’Connell for technical support, assistance and advice. We thank C. Alving and Z. Beck for designing the adjuvant.

## Funding

We acknowledge support from the U.S. Department of Defense, Defense Health Agency (Restoral FY20). This work was also partially executed through a cooperative agreement between the U.S. Department of Defense and the Henry M. Jackson Foundation for the Advancement of Military Medicine, Inc. (W81XWH-18-2-0040) and was supported in part by the US Army Medical Research and Development Command under Contract No. W81-XWH-18-C-0337. The views, opinions and/or findings are those of the authors and should not be construed to represent the positions, policy or decision of the U.S. Army or the Department of Defense. Research was conducted in compliance with the Animal Welfare Act and other federal statutes and regulations relating to animals and experiments involving animals and adheres to principles stated in the Guide for the Care and Use of Laboratory Animals, NRC Publication, 1996 edition.

## Author Contributions

H.A.D.K., M.G.J., S.V., N.L.M., K.M. and D.L.B. designed the study. I.E.N, A.A., C.M.C., C.S., K.K.P., H.H.H., R.E.C, P.V.T., W-H.C., R.S.S., A.H., E.J.M., C.E.P., W.C.C., M.C., A. Ahmed, K.M.W., M.D., I.S., J.B.C., K.G.L, V.D., S.M., K.A., R.C., S.J.K. D.P.P., N.K., V.R.P., Y.H., L.L.J., G.D.G. performed immunologic and virologic assays. H.A.E, A.C., M.G.L. led the clinical care of the animals. S.P.D., X.Z., E.K.D performed histopathology. K.M., M.G.J., P.V.T., W-H.C., R.S.S., A.H., E.J.M., C.E.P., W.C.C., and M.C. designed the immunogens. M.R., G.R.M., and A. Anderson designed and provided the adjuvant. H.A.D.K., M.G.J., C.S., J.A.H., M.F.A., S.P.D., P.T.S., D.D.H., M.S.D., M.R., G.D.D., S.A.P., N.L.M., K.M. and D.L.B. analyzed and interpreted the data. H.A.D.K., C.S., K.M. and D.L.B. wrote the paper with assistance from all coauthors.

## Competing Interests

M.G.J. and K.M. are named as inventors on International Patent Application No. WO/2021/21405 entitled “Vaccines against SARS-CoV-2 and other coronaviruses.” M.G.J. is named as an inventor on International Patent Application No. WO/2018/081318 entitled “Prefusion Coronavirus Spike Proteins and Their Use.” M.S.D. is a consultant for Inbios, Vir Biotechnology, NGM Biopharmaceuticals and Carnival Corporation and on the Scientific Advisory Boards of Moderna and Immunome. The Diamond laboratory has received funding support in sponsored research agreements from Moderna, Vir Biotechnology and Emergent BioSolutions. The other authors declare no competing interests.

## Materials & Correspondence

Correspondence and requests for materials should be addressed to K.M. and D.L.B.

