## Supplemental Figures and Tables for "Efficacy and breadth of adjuvanted SARS-CoV-2 receptor-binding domain nanoparticle vaccine in macaques"

**Supplemental Table 1. Nonhuman primate age, sex and weight distribution by group**

| ID | Sex | Age in Years | Weight (kg) | Group* | Immunogen |
|---|---|---|---|---|---|
| HS1608116 | F | 4.03 | 4.54 | 1-A | PBS |
| 171230 | F | 6.4 | 6.68 | 1-A | PBS |
| HS1605516 | F | 4.24 | 4.5 | 1-A | PBS |
| 180231 | M | 4.37 | 7.28 | 1-A | PBS |
| HS1610040 | F | 3.88 | 3.95 | 1-B | PBS |
| HS1606329 | F | 4.17 | 4.14 | 1-B | PBS |
| HS1606025 | M | 4.23 | 3.95 | 1-B | PBS |
| HS1606397 | M | 4.16 | 3.84 | 1-B | PBS |
| 171276 | F | 6.48 | 6.66 | 4-A | 50 µg RFN |
| HS1606381 | M | 4.16 | 4.22 | 4-A | 50 µg RFN |
| HS1606379 | M | 4.16 | 4.25 | 4-A | 50 µg RFN |
| HS1704246 | F | 3.35 | 4.16 | 4-B | 50 µg RFN |
| HS1607114 | F | 4.13 | 3.65 | 4-B | 50 µg RFN |
| HS1603140 | F | 4.46 | 3.4 | 4-B | 50 µg RFN |
| HS1603067 | M | 4.47 | 3.8 | 4-B | 50 µg RFN |
| HS1606371 | M | 4.16 | 3.94 | 4-B | 50 µg RFN |
| HS1610052 | F | 3.88 | 4.5 | 5-A | 5 µg RFN |
| 180654 | F | 5.46 | 4.16 | 5-A | 5 µg RFN |
| HS1606341 | M | 4.17 | 4.25 | 5-A | 5 µg RFN |
| 180288 | M | 4.46 | 8.44 | 5-A | 5 µg RFN |
| HS1704226 | F | 3.35 | 3.7 | 5-B | 5 µg RFN |
| HS1610030 | F | 3.88 | 4.1 | 5-B | 5 µg RFN |
| HS1606389 | M | 4.16 | 3.94 | 5-B | 5 µg RFN |

*A and B cohorts were sacrificed at day 14 and day 7 post-challenge, respectively

**Supplemental Table 2. Primers and probes for SARS-CoV-2 sgmRNA and total RNA viral load**

| Primer/Probe Name | Sequence 5'- 3' | Nucleotide Length |
|---|---|---|
| SARS-CoV-2 TAL E1 F | TCGTGGTATTCTTGCTAG | 18 |
| SARS-CoV-2 TAL E1 R | GAAGGTTTTACAAGACTCAC | 20 |
| SARS-CoV-2 TALE1 Probe | FAM -ACACTAGCCATCCTTACTGCG-BHQ1 | 21 |
| SARS-CoV-2 sg Leader | CGATCTCTTGTAGATCTGTTCTC | 23 |
| MS2 F | CTCTGAGAGCGGCTCTATTGG | 21 |
| MS2 R | GTTCCCTACAACGAGCCTAAATTC | 24 |
| MS2 Probe | JOE-TCAGACACGCGGTCCGCTATAACGAT- BHQ2 | 26 |

F = Forward Primer, R = Reverse Primer

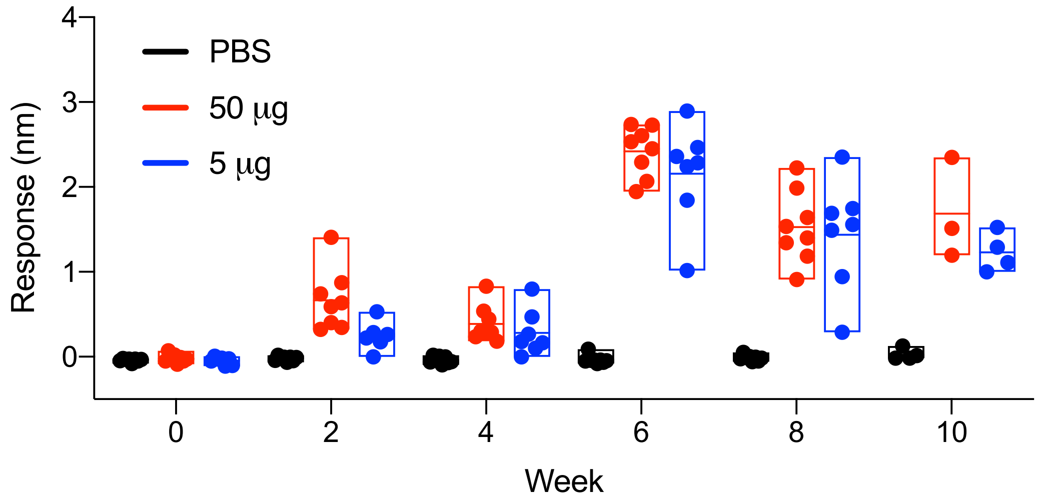


**Supplemental Figure 1. Binding antibody responses to SARS-CoV-2 RBD measured by biolayer interferometry.** SARS-CoV-2 RBD-specific binding antibody responses were assessed in macaque serum every two weeks following RFN immunization (weeks 0, 4) and challenge (week 8).

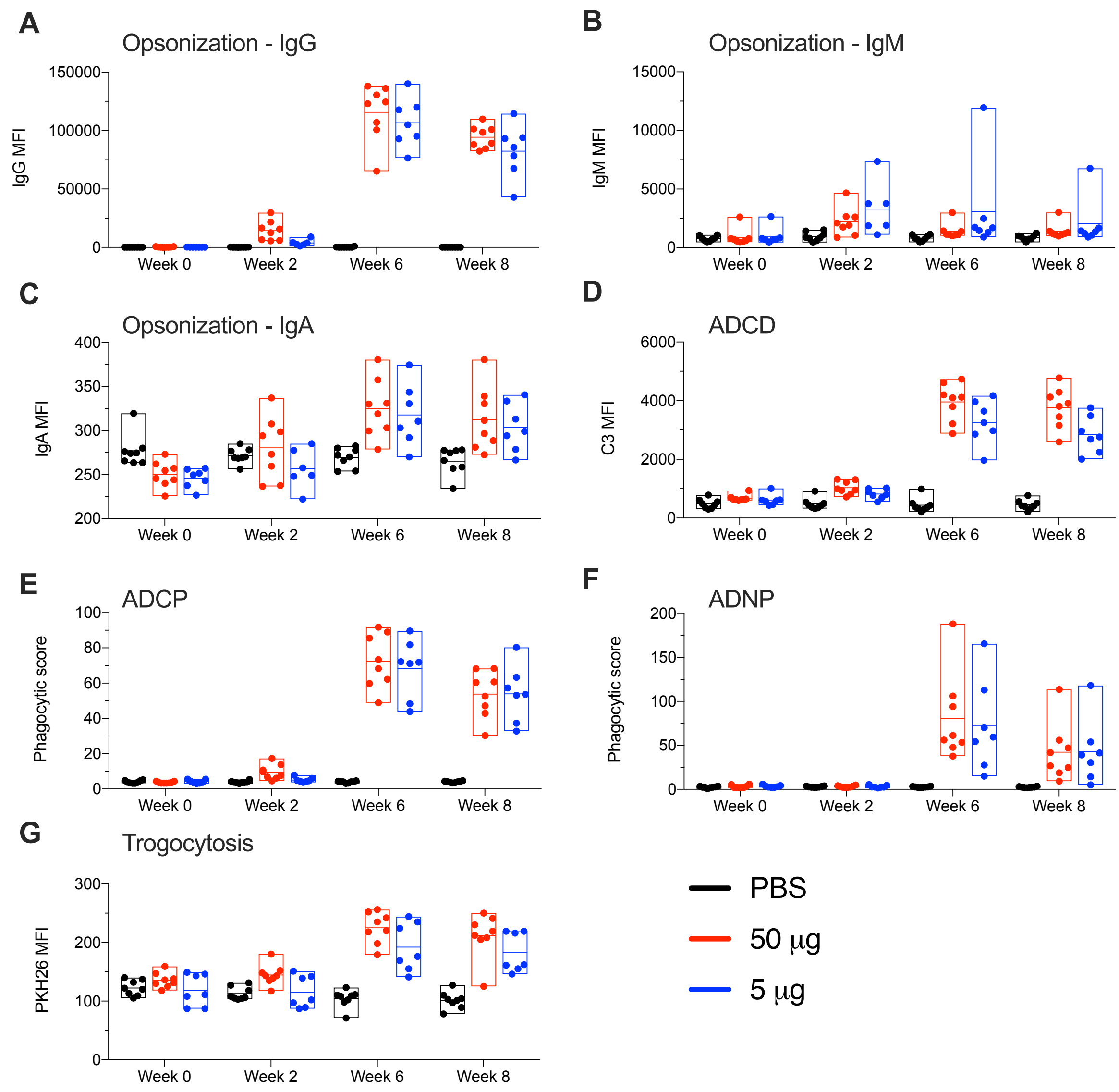


**Supplemental Figure 2. Fc-mediated effector antibody responses induced by vaccination with RFN.** SARS-CoV-2 (USA-WA1) S-specific plasma antibody effector activity was measured in RFN vaccinated macaques at the indicated study weeks. (*A-C*) Antibody-mediated cellular opsonization activity was measured using S-expressing cells incubated with diluted plasma followed by IgG (A), IgM (B), and IgA (*C*) staining detected by flow cytometry. (*D*) ADCD was measured on plasma-opsonized S-expressing cells. (*E,F*) ADCP and ADNP responses assessed by incubating spike-trimer-coated fluorescent beads with diluted plasma and culture with effector cells. (*G*) Trogocytosis was measured using plasma-opsonized S-expressing cells.

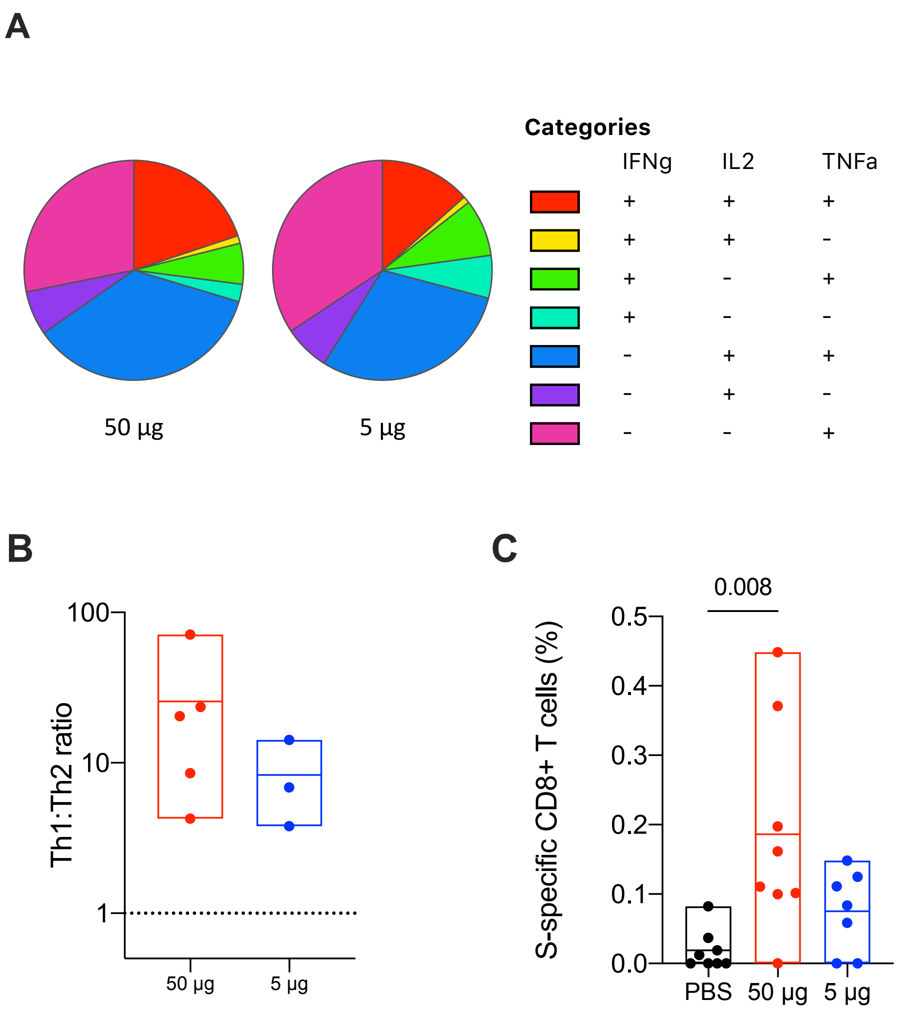


**Supplemental Figure 3. CD8+ T cell responses and CD4+ T helper response ratio and polyfunctionality.** SARS-CoV-2 (USA-WA1) S-specific T cell responses were assessed in PBMC of RFN vaccinated rhesus macaques four weeks after the last immunization by intracellular cytokine staining (*A*) S-specific CD4+ T cell Th1 cytokine polyfunctionality was assessed by Boolean combination gating of IFNγ, IL-2, and TNF expression. (*B*) The ratio of Th1 to Th2 S-specific memory CD4+ T cells in animals with positive Th2 responses. Dashed line indicates an equal proportion of Th1 and Th2 cells. (*C*) Memory CD8+ T cell responses were measured by stimulation with overlapping SARS-CoV-2 S peptides and IFNγ, IL-2 and TNF intracellular staining. Significance was assessed using a Kruskal-Wallis test followed by a Dunn's post-test.

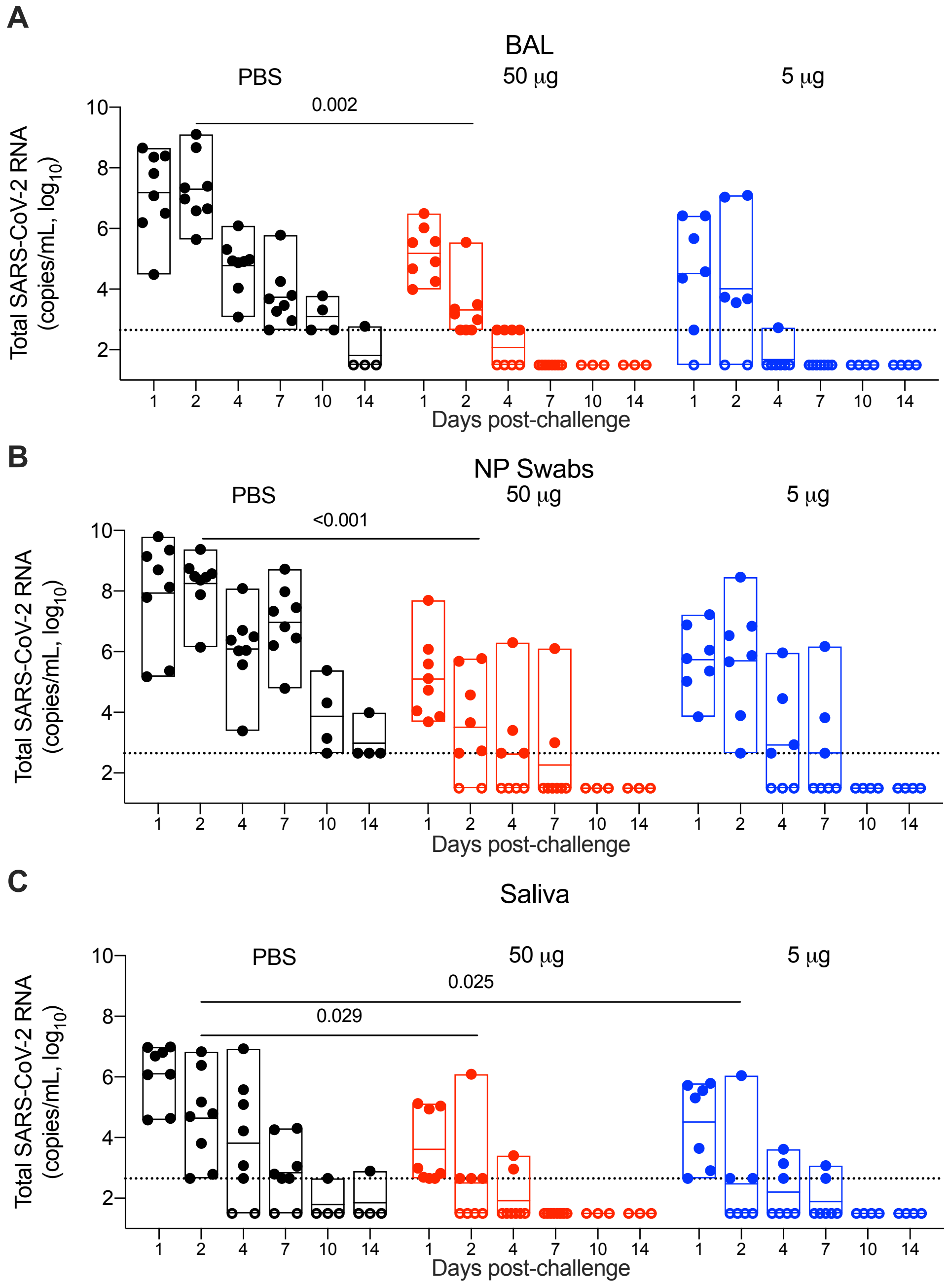


**Supplemental Figure 4. Total viral load in the airways following SARS-CoV-2 respiratory tract challenge.** Total SARS-CoV-2 RNA copies per milliliter were measured in bronchoalveolar lavage fluid (*A*), nasopharyngeal swabs (*B*) and saliva (*C*) of vaccinated and control animals for two weeks following intranasal and intratracheal SARS-CoV-2 (USA-WA1/2020) challenge. Specimens were collected 1, 2, 4, 7, 10 and 14 days post-challenge (N=7-8 per group for days 1-7; N=3-4 days 10 and 14). Dotted lines demarcate assay lower limit of linear performance range (log10 of 2.65 corresponding to 450 copies/mL); positive values below this limit are plotted as 450 copies/ml. Open symbols represent animals with viral loads below the limit of detection of the assay. Box plot horizontal lines indicate the mean; top and bottom reflect the minimum and maximum. Significant differences between control and vaccinated animals at day 2 post-challenge are indicated. Significance was assessed using a Kruskal-Wallis test followed by a Dunn's post-test.

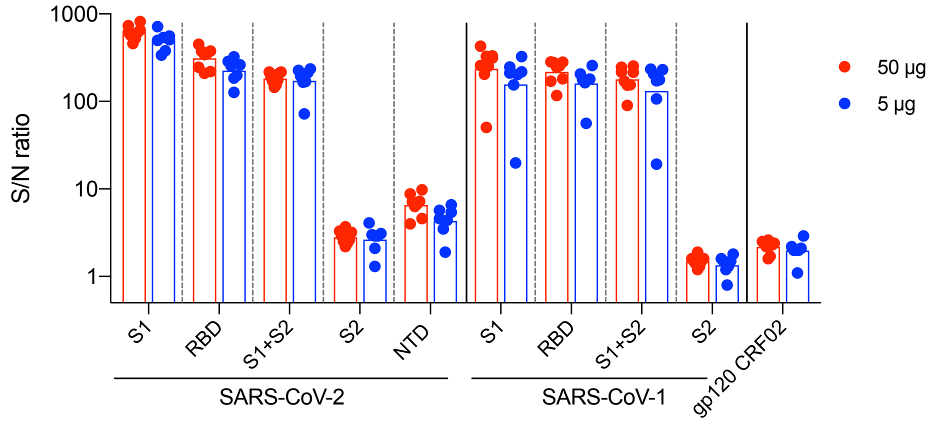


**Supplemental Figure 5. Binding antibody responses to SARS-CoV-2 and SARS-CoV-1 antigens measured by Luminex.** RFN vaccinated macaque plasma collected two weeks after the last immunization was evaluated for binding to SARS-CoV-2 and SARS-CoV-1 S1 and S2 subunits, RBD, and N-terminal domain (NTD) using a multiplex Luminex assay. Mean fluorescence intensity (MFI) data were divided by pre-immunization background MFI to obtain a signal to noise (S/N) ratio of vaccine-elicited binding antibody response magnitude for each sample. Antibody binding to HIV gp120 circulating recombinant form (CRF) 02 was used as a negative control antigen. Bars indicate the geometric mean.

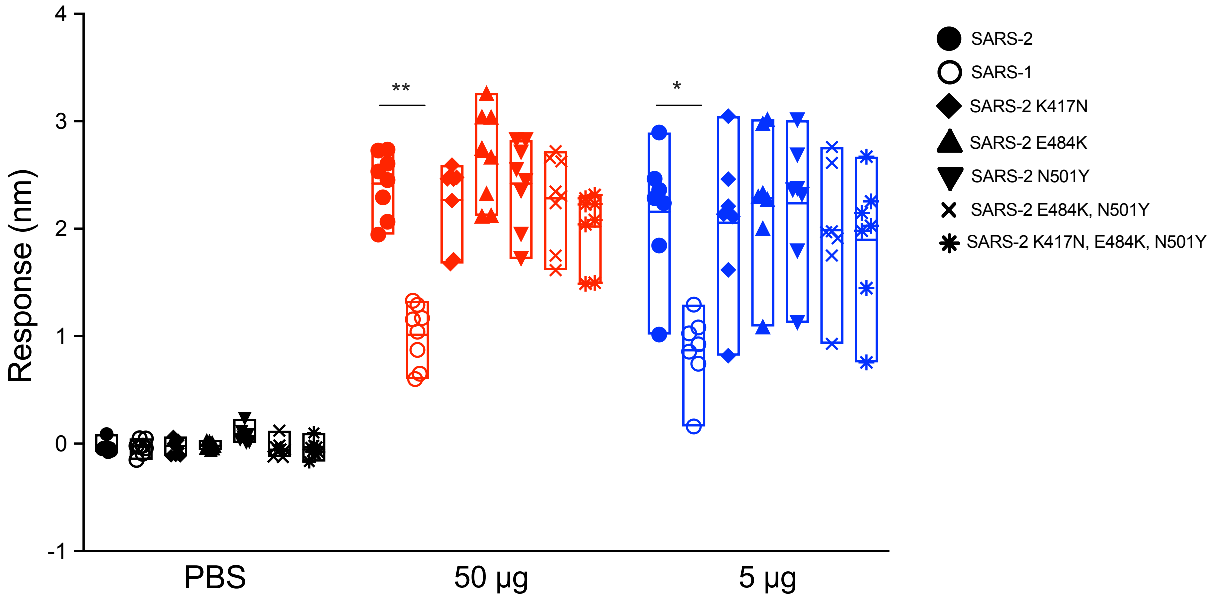


**Supplemental Figure 6. Binding antibody responses to SARS-CoV-1, SARS-CoV-2 wild-type and SARS-CoV-2 mutant RBD.** Serum RBD-specific antibody responses were assessed by biolayer interferometry two weeks after last RFN vaccination**.** SARS-CoV-2 RBD variant forms produced by site-directed mutagenesis were used as antigens. Significant differences relative to SARS CoV-2 wild-type binding assessed using a Kruskal-Wallis test followed by a Dunn's post-test is indicated (* <0.05; ** < 0.01).

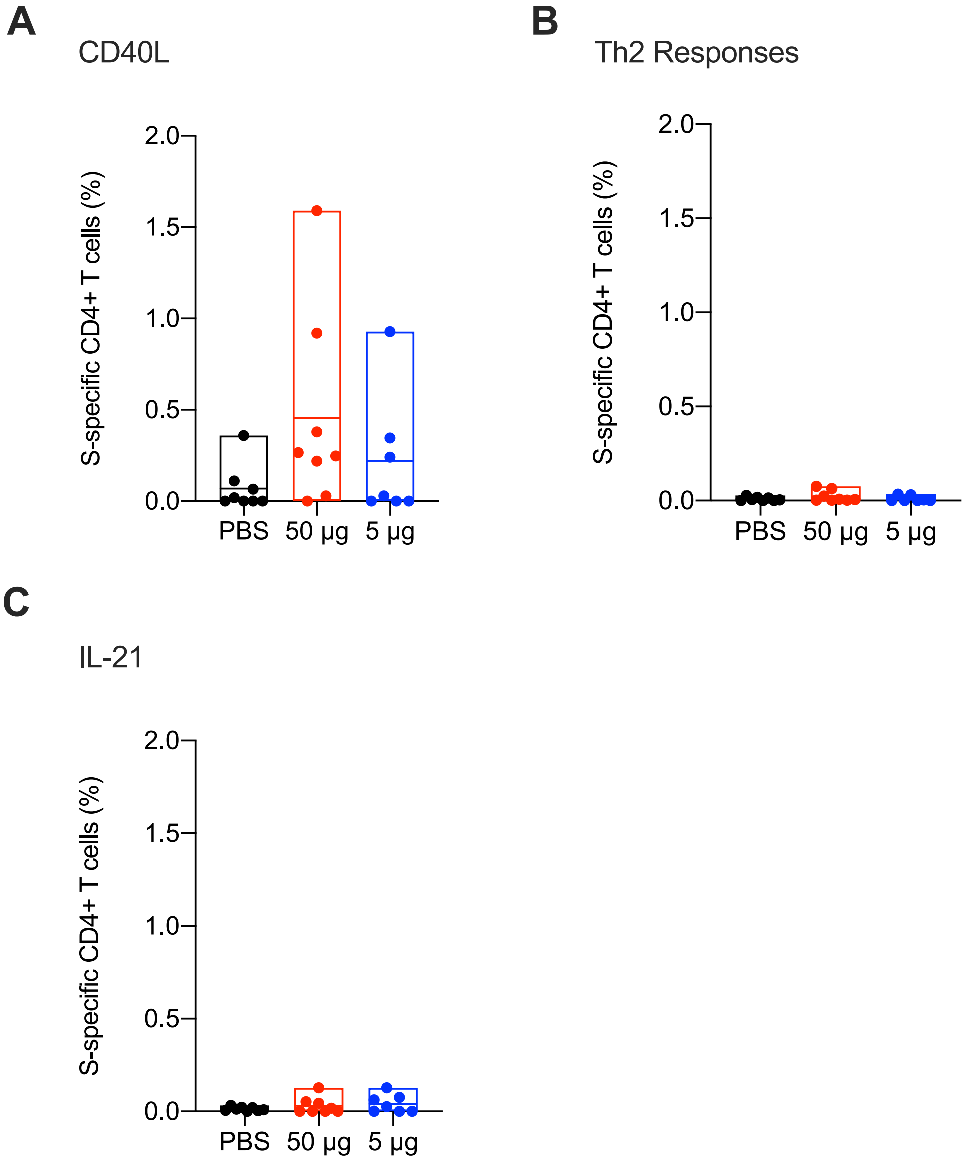


**Supplemental Figure 7. Cross-reactive CD4+ T cell responses against SARS-CoV-1.** Antigen-specific T cell responses were assessed in RFN vaccinated and control macaques by SARS-CoV-1 S peptide pool stimulation of PBMC collected two weeks after the last vaccination followed by ICS. The frequency of S-specific memory CD4+ T cells expressing the indicated marker(s) is shown for (*A*) CD40L, (*B*) Th2 cytokines (IL-4 and IL-13) and (*C*) IL-21. Boolean combinations of cytokine positive memory CD4+ T cells were summed. Significance was assessed using a Kruskal-Wallis test followed by a Dunn's post-test.
